# “IGF2BP1 phosphorylation regulates ribonucleoprotein condensate formation by impairing low-affinity protein and RNA interactions”

**DOI:** 10.1101/2023.12.05.570121

**Authors:** Harald Hornegger, Adnan Muratovic, Aleksandra Anisimova, Benjamin Burgeois, Elena Spinetti, Roberto Covino, Tobias Madl, G Elif Karagöz

## Abstract

The insulin-like growth factor 2 mRNA binding protein (IGF2BP1) is a conserved RNA-binding protein that regulates RNA stability, localization, and translation. IGF2BP1 is part of various ribonucleoprotein (RNP) condensates regulating RNA outputs. However, the mechanism that regulates its assembly into condensates remains unknown. Here we found, using proteomics, that IGF2BP1 phosphorylation at S181 in a disordered linker is regulated in a stress-dependent manner. Phosphomimetic mutations in two disordered linkers, S181E and Y396E, modulated RNP condensate formation by IGF2BP1 without impacting its binding affinity for RNA. Intriguingly, the S181E mutant, which lies in linker 1, impaired IGF2BP1 condensate formation *in vitro* and in cells, whereas a Y396E mutant in the second linker increased condensate size and dynamics. Structural approaches showed that the first linker binds RNAs nonspecifically through its RGG/RG motif, an interaction weakened in the S181E mutant. Notably, linker 2 interacts with IGF2BP1’s folded domains and these interactions were partially impaired in the Y396E mutant. Our data reveal how phosphorylation modulates low affinity interaction networks in disordered linkers to regulate RNP condensate formation.

## Introduction

RNA-binding proteins (RBPs) play important roles in post-transcriptional control of RNA ^1–6^. IGF2BPs are a conserved family of RBPs that regulate RNA localization, translation and stability ^7–11^. There are three IGF2BP paralogs (IGF2BP1-3) in mammals. Discovered in chicken embryos, IGF2BP1 was the founding member of the IGF2BP family ^12, 13^. IGF2BP1 is highly conserved in sequence and function across species (**Fig. Supp. 1A)**. It is highly expressed during mid to late embryogenesis and its expression decreases in adult tissues. In line with embryonic functions, *Igf2bp1* knockout mice show developmental abnormalities ^14^. However, IGF2BP1 expression is not restricted to early development and it is detected later in differentiated gonads and the kidneys. Consistent with post-developmental functions, loss of IGF2BP1 in intestinal epithelial cells impairs intestinal homeostasis in adults^15, 16^. IGF2BP1 is highly expressed in various tumors and its overexpression correlates with tumor aggressiveness ^9, 17^. Importantly, IGF2BP1 depletion impairs tumor growth, indicating that inhibition may have therapeutic potential in cancer cells ^18,19^. This link to disease underlines the importance of obtaining a mechanistic understanding of how IGF2BP1 exerts its function.

IGF2BP1 is a canonical multi-domain RBP, which contains six RNA-binding domains: two RNA recognition motif (RRM) domains and four hnRNP K homology (KH) domains that are linked by two intrinsically disordered regions (**Fig. 1A**). The KH domains are arranged into pseudodimers (KH1-2, and KH3-4). RNA recognition by IGF2BP1 is mediated by the KH domains, which interact with single-stranded RNAs through 4 nucleotide long recognition motifs ^20^. In contrast, the RRM domains provide little specificity and promiscuously recognize dinucleotide sequences, as shown for the IGF2BP3 paralog ^21, 22^. These multivalent interactions increase the specificity and affinity of IGF2BPs for substrate RNAs ^20, 21, 23^.

**Figure 1.**
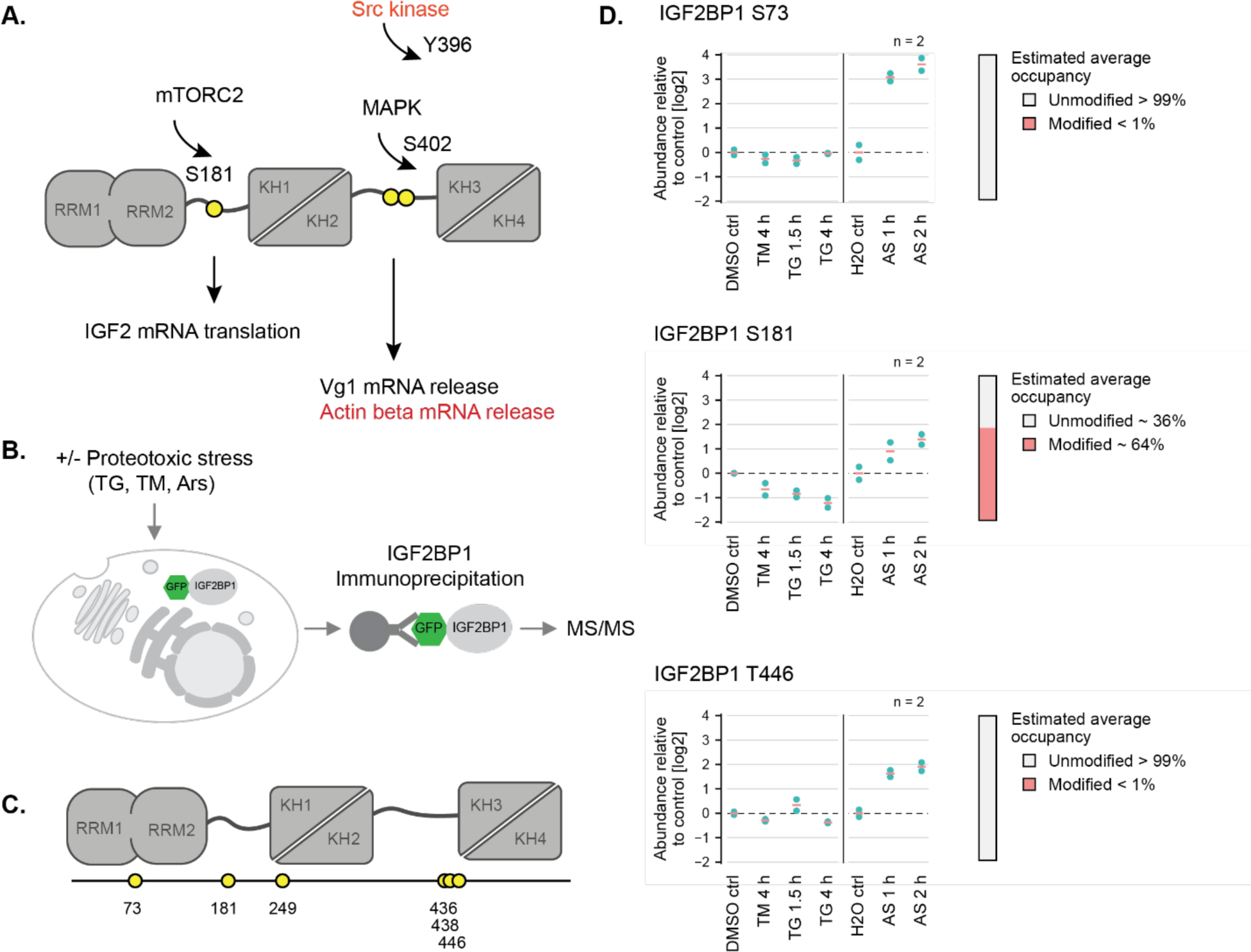
**(A)** Schematic depiction of IGF2BP1 domain architecture. IGF2BP1 consists of 6 RNA binding domains: two RRM domains and four KH domains, which are arranged as pseudo-dimers connected via two linkers. The well-studied phosphorylation sites (S181, Y396, S402), the respective kinases and the effect of the phosphorylation are also depicted. **(B)** Schematic overview of the workflow for mass spectrometry experiments to determine the stress-regulated phosphorylation sites in IGF2BP1. **(C)** Representation of IGF2BP1 phosphorylation sites identified by MS analyses. **(D)** Relative abundance of the indicated IGF2BP1 phosphorylation sites in cells exposed to various forms of proteotoxic stress compared to the control conditions. Tunicamycin (TM) and thapsigargin (TG) induces ER stress, whereas sodium arsenite (AS) leads to oxidative stress. The time-points on the bottom indicate length of exposure to the stress-inducing drug.

Genome-wide cross-linking and immunoprecipitation (CLIP) studies identified a large number of IGF2BP1 targets, suggesting roles in cell growth, migration, synaptic plasticity in healthy tissues, as well as tumor growth and metastasis in cancer cells ^9, 17, 24–27^. These data also revealed that IGF2BP1 binds to the coding regions, 5’-untranslated regions (UTRs), and 3’-UTRs of target RNAs, with the highest number of binding sites residing in 3’-UTRs ^28, 29^. Since the binding sites for IGF2BP1 and microRNAs overlap, it was proposed that IGF2BP1 can stabilize RNAs by competing with the microRNA binding sites ^19^. IGF2BP1 also binds to and stabilizes N^6^ methyl adenosine-modified RNAs during heat shock stress ^8^. Although IGF2BP1 has been proposed to stabilize RNAs, binding to a subset of its target RNAs correlates with destabilization^30^. However, what regulates these distinct functional outputs remains largely unknown.

IGF2BP1 assembles into various ribonucleoprotein granules (RNP) to regulate RNA fate. In neurons, IGF2BP1 is part of transport granules, which transport select mRNAs from soma to neurites to regulate site-specific protein synthesis. During cellular stress, IGF2BP1 is sequestered into stress granules that have been proposed to protect mRNAs from degradation until translation resumes. Intriguingly, IGF2BP1 also localizes to P-bodies, which are sites of RNA recapping and degradation. Yet, the regulation of IGF2BP1 assembly into RNP granules with opposite functions is not well understood.

One well-defined mechanism that regulates IGF2BP1 function is through its phosphorylation. In the best-studied example, phosphorylation of IGF2BP1 controls its binding to the ß-actin-encoding *ACTB* mRNA, providing a regulatory switch to allow for spatial control of *ACTB* mRNA translation ^7^. IGF2BP1 binds to the 3’-UTR of the *ACTB* mRNA and prevents its translation. Phosphorylation of IGF2BP1 at Y396 by the Src kinase, which is localized to the leading edge of the cell or axons, releases IGF2BP1 from *ACTB* mRNAs, allowing their translation at those sites ^7^. In contrast, the phosphorylation of IGF2BP1 at S181 was suggested to enhance its binding to the 5’-UTR *of IGF2* mRNA, thereby increasing *IGF2* mRNA translation ^31, 32^. Importantly, S181 phosphorylation regulates stabilization of IGF2BP1 target RNAs and impacts dendritic branching in hippocampal neurons underlining its functional importance ^33,34^. Yet, although IGF2BP1 phosphorylation at distinct sites has been suggested to impact RNA binding, the mechanistic details of IGF2BP1 regulation by phosphorylation remains only partially understood. Moreover, whether IGF2BP1 is regulated by other phosphorylation events has yet to be examined.

Here, we map stress-regulated phosphorylation sites in IGF2BP1 by targeted mass spectrometry analyses to uncover mechanisms that control IGF2BP1 outputs. Using *in vitro* reconstitution, biochemistry, and structural methods, we dissect how IGF2BP1 phosphorylation in its disordered linker regions regulates function. We show that phosphorylation of the disordered linkers regulates the propensity of IGF2BP1 to form RNP granules *in vitro* and in cells by modulating low affinity interaction networks. Our data reveal how disordered regions provide highly tunable regulation of RNP condensate formation through a single phosphorylation event.

## Results

### IGF2BP1 is phosphorylated during proteotoxic stress

IGF2BP1 stabilizes a subset of RNAs during proteotoxic stress and we therefore tested whether IGF2BP1 is regulated through phosphorylation under those conditions ^8, 35^. To this end, we mapped phosphorylation sites in IGF2BP1 by mass spectrometry (MS) in mammalian cells under control conditions and under conditions where the cells were exposed to proteotoxic stress (**Fig. 1B**). To compare IGF2BP1 phosphorylation sites under various forms of proteotoxic stress, we exposed the cells to oxidative stress using sodium arsenite (1 or 2 hrs) or endoplasmic reticulum (ER) stress using tunicamycin (4 hrs) or thapsigargin (1.5 and 4 hrs).

To increase specificity and stringency in our analyses, we enriched for IGF2BP1 by immunoprecipitation from HEK293 cells engineered by CRISPR/Cas9 gene editing to express GFP-tagged IGF2BP1 using split-GFP technology ^36^. We obtained a sequence coverage with identified peptides covering 78.7% of the IGF2BP1 amino acid sequence. Peptides covering the disordered linker 2, spanning from amino acids 347 to 423, were not detected in the MS analyses. This likely resulted from the low complexity nature of this region, which may be inaccessible to tryptic digestion to generate MS-compatible peptides. Thus, we were not able to investigate the phosphorylation status in the linker 2.

MS analyses identified several IGF2BP1-derived phosphopeptides whose levels increased or decreased in a stress-dependent manner (**Fig. 1C-D, Fig. Supp. 1B**). The identified phosphopeptides mapped to the RRM1 domain (aa S73), disordered linker 1 (aa S181), KH1 (aa T249), KH2 and KH3 domains (aa S436, S438, T446) (**Fig. 1C)**. The ratio of phosphorylated to unmodified peptides was relatively low (< 1% of total protein) for most of the identified phosphopeptides (S73, T249, S436, S438, T446). The most prominent phosphorylation site we identified mapped to S181 (aa 176-QPRQG**S**PVAAGA-187 and > 64% of total protein), which increased by around two-fold during oxidative stress induced by arsenite treatment. In contrast, S181 phosphorylation decreased two-fold when cells were treated with ER stress-inducing drugs, indicating that this phosphorylation event depends on stress type. Multiple amino acid sequence alignments revealed that S181 is highly conserved from fish to mammals, suggesting its functional importance (**Fig. Supp. 1A)**. This site in has previously been proposed to be phosphorylated in all three IGF2BP paralogs by mTORC2 ^31, 32^. Notably, apart from mTORC2, motif prediction based on recent work^37^ indicated that S181 might be phosphorylated by members of the CMGC kinase family (i.e. SRPK2 and DYRK3), suggesting that other kinases might be involved in this regulation. Altogether, we found that IGF2BP1 is phosphorylated at multiple sites in a stress-dependent manner.

### Phosphomimetic mutants do not impact IGF2BP1 interaction with RNA

Since S181 was the most prominent stress-regulated phosphorylation site we identified, we went on to dissect how it regulates IGF2BP1 function. We first tested whether S181 phosphorylation regulates IGF2BP1 interaction with RNAs. Interestingly, whereas phosphorylation of S181 in linker 1 is proposed to increase its binding to RNAs, while phosphorylation of Y396 in disordered linker 2 was proposed to decrease it (**Fig. 1A)** ^7, 31, 32^. However, how phosphorylation of IGF2BP1 at the disordered linkers regulates its function remained only partially understood. To test whether, as proposed, linker phosphorylation impacts IGF2BP1 interaction with RNA, we used *in vitro* assays to measure binding of wild-type IGF2BP1 and its phosphomimetic mutants (IGF2BP1 S181E and Y396E) to model RNAs. For these experiments, we selected two IGF2BP1 target mRNAs based on published CLIP data sets, the unfolded protein response transcription factor *XBP1* and the translation initiation factor *EIF2A*. CLIP data showed that IGF2BP1 crosslinks to a distinct region in the 3’-UTR of the *XBP1* (**Fig. Supp. 2A, B, Table 1)** and that *EIF2A* mRNA is enriched in several predicted IGF2BP1-binding motifs ^29^ (**Table 1**).

IGF2BP binds RNA targets through a combinatorial code recognized by all six RNA-binding domains ^21, 23^. This interaction involves recognition of a cluster of distinct and regularly spaced RNA elements covering a ∼100 nucleotide-long target RNA region ^21^. We *in vitro* transcribed approximately 200 nt-long regions from the *XBP1* and *EIF2A* RNAs (**Table1**) and tested their binding to IGF2BP1 using Electrophoretic Mobility Shift Assays (EMSA) assays. IGF2BP1 bound to the 3’-UTR of *XBP1* and *EIF2A* with similar affinities (*XBP1*, K_D_= 41.0 nM and *EIF2A* at K_D_=48.2 nM) (**Fig. 2A, Fig. Supp. 2C-F, Table 2)**. We wondered whether IGF2BP1 phosphorylation at the linkers would impact the combinatorial recognition of RNA motifs by the individual RNA-binding domains, thereby influencing IGF2BP1’s binding to RNAs. Phosphomimetic IGF2BP1mutants (S181E and Y396E) bound to the XBP1 201 nt RNA with a slightly higher affinity than wild-type IGF2BP1 (**Fig. 2B, C, Fig. Supp. 2C, Table 2)** (K_D_, S181E= 17.1 nM, K_D_, Y396E= 22.9 nM). A similar affinity and effect of the phosphomimetic mutants was observed for their interaction with the EIF2A 200 nt RNA (**Fig. Supp. 2D-F, Table 2)** (K_D_, wild-type: 48.2 nM, K_D_, S181E: 35.4 nM, K_D_, Y396E: 40.2 nM).

**Figure 2.**
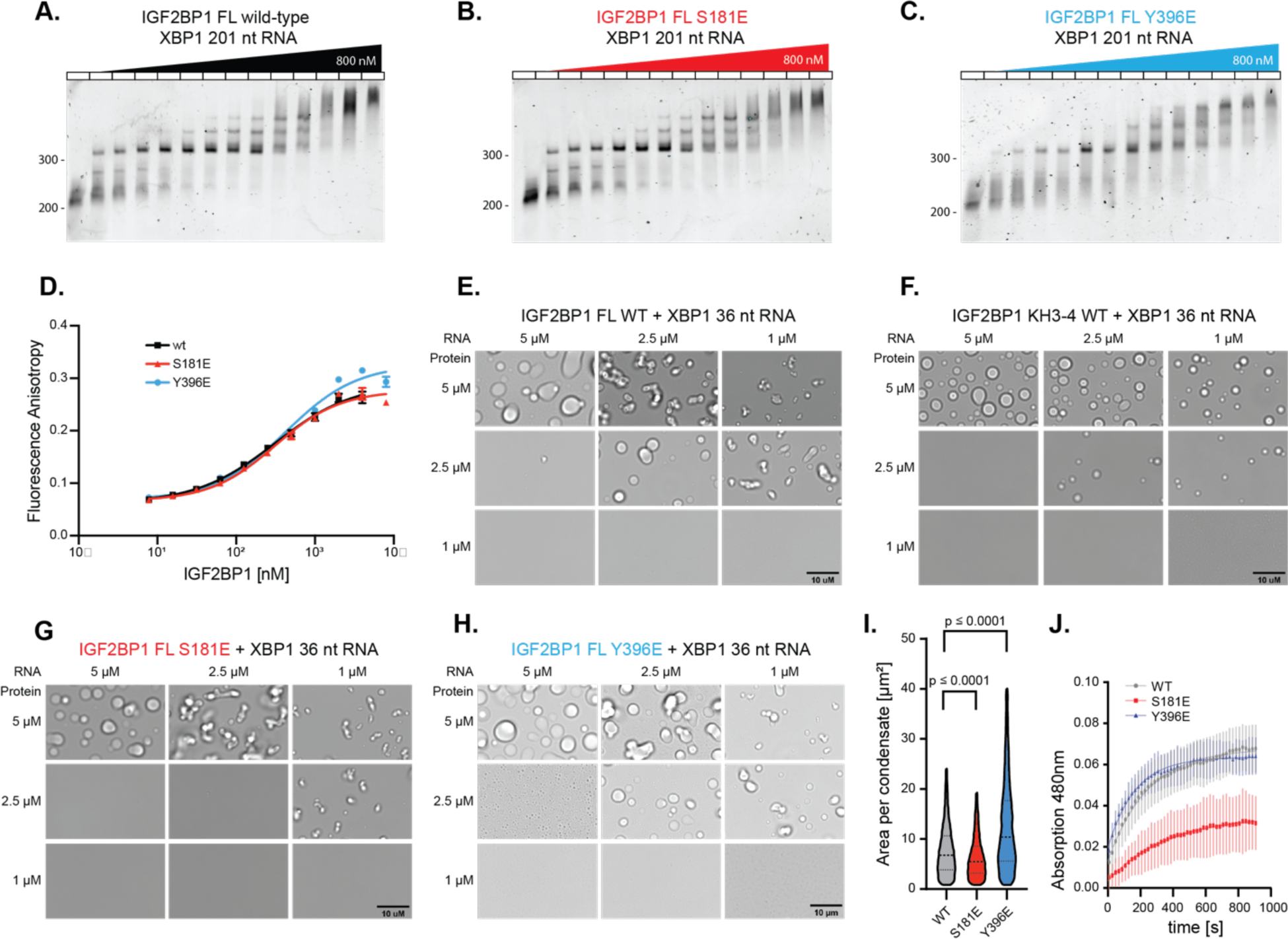
**(A)** Electrophoretic Mobility Shift Assay (EMSA) with XBP1 201 nt RNA to assess the binding of wild-type IGF2BP1 at concentrations ranging from 0 to 800 nM. **(B)** EMSA assays with IGF2BP1 S181E mutant under the same conditions as Fig.2A. **(C)** EMSA assays with IGF2BP1 Y396E mutant under the same conditions as Fig.2A. **(D)** Fluorescence anisotropy experiments to assess binding of wild-type IGF2BP1 (black), S181E (red) and Y396E (blue) to 5’-fluorescein labeled XBP1 36 nt RNA. Error bars represent the standard deviation. X-axis is represented in log-scale. **(E)** RNP granule formation of wild-type IGF2BP1 in the presence of XBP1 36 nt RNA after 90 min of incubation. The protein and RNA concentrations are indicated in the figures. **(F)** RNP granule formation assays IGF2BP1 KH3-4 pseudodimers **(G)** IGF2BP1 S181E and **(H)** IGF2BP1 Y396E mutants in the presence of XBP1 36 nt RNA after 90 min of incubation. Scale bar is 10 µm. **(I)** Violin plot of the area per condensate after 90 min incubation with 5µM IGF2BP1 full-length wild-type (black), S181E (red) and Y396E (sblue) with 5 µM XBP1 36 nt RNA. The condensates were quantified by fluorescence microscopy using mCherry-IGF2BP1 signal of the respective mutant. **(J)** RNP condensate formation kinetics monitored by a turbidity assay by measuring the light scattering at 480 nm of condensates formed by IGF2BP1 full-length wild-type (grey), S181E (red) and Y396E (blue) in the presence of XBP1 36 nt RNA over the course of 15 min. Error bars represent the standard deviation. Fitted curve represents a one-phase association equation. n = 10

As a complementary quantitative approach, we set up fluorescence anisotropy assays to measure the affinity of IGF2BP1 for shorter RNAs. By truncating the 3’-UTR of the *XBP1* mRNA, we identified a 36 nt-long RNA that is composed of two predicted IGF2BP1 recognition motifs (**Fig. Supp. 2B, Table 1)**. Fluorescence anisotropy assays showed that 5’-fluorescein-tagged XBP1 36 nt RNA bound to the wild-type IGF2BP1 with an affinity that was an order of magnitude lower than the 201 nt-long version (**Fig. 2D**, **Table 2**, K_D_, wild-type: 311.7 nM). We speculate that the lower affinity results from a reduced number of binding sites in the RNA, resulting in the decreased potential for combinatorial binding by the RNA-binding domains. IGF2BP1 phosphomimetic mutants S181E and Y396E bound to XBP1 36 nt RNA at a comparable affinity (**Fig. 2D**, **Table 2**, K_D_, S181E: 310.1nM, K_D,_ Y396E: 423.3 nM) to the wild-type IGF2BP1, indicating that the phosphomimetic mutants do not significantly impact IGF2BP1s interaction with model RNAs. These findings are consistent with previous data indicating that canonical folded RNA-binding domains in IGF2BP1 drive its interaction with target RNAs.

### IGF2BP1 forms RNA-mediated RNP granules in vitro

IGF2BP1 function has been associated with its assembly into RNP granules ^38–40^. Therefore, we investigated whether IGF2BP1 phosphorylation impacts formation of IGF2BP1 condensates. To test this possibility, we aimed to reconstitute RNP granules formed by IGF2BP1 and RNAs. Many RNP granules form through liquid-liquid phase separation (LLPS) driven by multivalent interaction between RBPs with RNAs. To allow multivalent binding of IGF2BP1 to RNAs, we used the XBP1 36 nt RNA, which contains two predicted IGF2BP1-binding motifs (**Fig. Supp. 2B, Table 1**). We incubated IGF2BP1 with this RNA at different concentrations and stoichiometry and monitored whether they form RNP condensates visible as droplets by bright-field microscopy (**Fig. 2E)**. Under physiological pH and salt conditions, 2.5 µM IGF2BP1 and 1 µM XBP1 36 nt RNA readily formed RNP condensates (**Fig. 2E, Fig. Supp. 3A, B**). Likewise, IGF2BP1 formed condensates with another model RNA (**Fig. Supp. 3C)**. Increasing full-length IGF2BP1 and/or RNA concentration resulted in formation of larger condensates, whereas excess RNA with higher than 1-fold molar ratio abrogates condensate formation. These data indicated that under these conditions, RNAs do not form homotypic contacts that contribute to condensation. Instead, they act as monomers competing for IGF2BP1 binding sites.

We next mapped which RNA-binding domains in IGF2BP1 contribute to formation of RNP condensates. To assess this, we first measured the affinity of IGF2BP1’s individual domains for model RNAs. We found that KH3-4 dimers bound to a model *ACTB*-derived RNA (ACTB 28 nt) ^41^ and the XBP1 36 nt RNA (**Table 1**) at around 1.5 µM affinity (**Fig. Supp. 3D, E, Table 2)**. Instead, KH1-2 bound to the same RNAs with an affinity of > 15 µM. These data are consistent with earlier work indicating that KH3-4 domains bind to RNA with the highest affinity ^42^. Introducing GEEG mutations, which impede the RNA interaction of the respective KH domain ^43, 44,43^, into the RNA-binding motif in the KH3 domain in KH3-4 dimers decreased binding affinity to XBP1 36 nt RNA by 10-fold (**Fig. Supp. 3F, Table 2**, K_D_=16.0 µM). In contrast, the KH3-4 mutant in which the KH4 binding site is mutated, bound to RNA with a similar affinity as the wild-type KH3-4 dimers (K_D_=2.0 µM). These data argue that KH3 provides the major RNA binding site since the affinity did not change compared to the wild-type KH3-4 construct.

A KH1-4 construct lacking the RRM1-2 dimers and linker 1 bound to the ACTB and XBP1 36 nt RNA with similar affinity as the full-length IGF2BP1 (**Fig. Supp. 3D, Table 2**, ACTB: K_D_= 245.4 nM, XBP1: K_D_= 204.0 nM). These data suggest that the KH1-2 and KH3-4 domains drive the avidity effect for short RNAs with two binding sites. To characterize this further, we mutated the KH3 and KH4 RNA-binding motifs to GEEG in the full-length IGF2BP1. Both the EMSA assays and fluorescent anisotropy experiments showed that full-length IGF2BP1 KH3-4 GEEG double mutant bound to RNA with similar affinity as the wild-type IGF2BP1 (**Fig. Supp. 3G,H, Table 2**, K_D_ mutant: 185.8 nM) and that simultaneous binding of RRM1-2 and KH1-2 dimers to RNA can also benefit from the avidity effect. Interestingly, the EMSA assays performed with the XBP1 201 nt RNA showed that compared to wild-type IGF2BP1, the IGF2BP1 KH3-4 GEEG double mutant displayed differences in the high molecular weight assemblies formed at higher protein concentrations (> 250 nM, **Fig. Supp. 3H**). From these data, we concluded that the impaired RNA-binding of KH3-4 pseudodimers results in a different mode of RNA recognition by the mutant.

In line with the fluorescence anisotropy experiments, KH3-4 domains alone formed condensates in the presence of RNA (**Fig. 2F**). Importantly, impairing RNA binding to either KH3 or KH4 domains through GEEG mutations abolished condensate formation (**Fig. Supp. 4A, B**). These mutants lacked the ability to form multivalent interactions required for condensate formation. Together with the fluorescence anisotropy assays, these data revealed that KH3-4 domains are necessary and sufficient to build the multivalency that drives IGF2BP1 RNP condensate formation. Consistent with these results, bright-field microscopy analysis showed that the full-length IGF2BP1 KH3-4 GEEG mutant did not form condensates (**Fig. Supp 4C**). These data indicated that even though this mutant binds to RNA with high affinity (**Fig. Supp. 3G, H, Table 2**, K_D_:185.8 nM), the low RNA-binding affinity of the individual KH1-2 (**Fig. Supp. 3D, E, Table 2**) and RRM1-2 domains does not allow the formation of multivalent interactions that are essential for condensate formation under those conditions. Supporting this, KH1-2 domains alone did not form condensates under conditions where KH3-4 formed droplets (**Fig. Supp. 4D**). Importantly, a model RNA with a single IGF2BP1-binding motif (XBP1 10 nt RNA) did not mediate condensate formation when incubated with KH1-4, validating that multiple binding-sites in both the RNA and protein are required for condensation (**Fig. Supp 4E**). Remarkably, KH1-4 formed mesh-like networks upon incubation with the XBP1 36 nt RNA at conditions that lead to the formation droplets with IGF2BP1 full-length wild-type (**Fig. Supp. 4F**). These data suggested that promiscuous RNA interactions by the RRM1-2 domains increases the dynamics of IGF2BP1-RNA interactions. In the presence of 250 mM NaCl, KH1-4 formed condensates similar to full length IGF2BP1 (**Fig. Supp. 4G)**. We speculate that presence of high concentration of salt weakens the interaction of KH1-4 with RNA increasing their binding dynamics in the condensates. In summary, our data revealed that IGF2BP1 forms condensates driven by binding of the KH3-4 pseudodimers to RNAs. We speculate that once the condensates are formed, due to the high protein and RNA concentration in the condensed phase, the RRM1-2 and KH1-2 pseudodimers can form additional contacts with RNA.

### IGF2BP1 phosphomimetic mutants impact formation and dynamics of IGF2BP1 RNP granules

After establishing that IGF2BP1 assembled into condensates together with RNA, we next tested whether phosphorylation of IGF2BP1 linker regions impacts formation of IGF2BP1 RNP condensates. To this end, we used fluorescence microscopy to quantify the size and area of the condensates at 90 minutes after condensate formation for wild-type IGF2BP1 and its phosphomimetic mutants. Quantification of the IGF2BP1-RNA condensates (**Fig. Supp. 3A**, see materials and methods) revealed that the size and the total area of condensates formed by the S181E mutant was smaller compared to the wild-type IGF2BP1, confirming that the IGF2BP1 S181E mutant is impaired in condensate formation (**Fig. 2G, I, Fig. Supp 4I-L, Table 3**, median area per condensate: wild-type: 7.0 µm, S181E: 5.7 µm, mean total area: wild-type: 7753 µm², S181E: 4691 µm², at 5 µM protein and RNA concentration). While IGF2BP1 S181E forms almost no condensates at 2.5 µM when incubated with stoichiometric amounts of XBP1 36 nt RNA (**Fig 2G**), the presence of 5% mCherry-labeled construct leads to the formation of small condensates (**Fig. Supp. 4I-L**). We observed that the mCherry-tag enhances the phase separation propensity of IGF2BP1. This effect was prominent when 2.5 µM IGF2BP1 full-length S181E was exposed to 2.5 µM XBP1 36 nt RNA (**Fig. Supp. 4J),** likely because the saturating concentration of this protein is very close to 2.5 µM. Thus, we used sub-stochiometric amounts of mCherry-labeled IGF2BP1 to quantify the condensate area. Intriguingly, in contrast to the S181E mutant, the Y396E mutant formed larger condensates with a larger total area under the same experimental conditions (**Fig. 2H, I, Fig. Supp 4I-L, Table 3**, median area per droplet: Y396E 10.8 µm, mean total area: Y396E 9298 µm², at 5 µM protein and RNA concentration). Similarly, incubation of XBP1 36 nt RNA with KH1-4 Y396E mutant led to formation of condensates with more regular droplet-like shapes compared to the condensates formed by the wild-type KH1-4 under the same conditions (250 mM NaCl). These data confirmed that Y396E mutation impacts condensate formation and this effect does not depend on RRM1-2 domains in IGF2BP1 (**Fig. Supp. 4H**).

We next used turbidity assays to monitor the kinetics of condensate formation following the addition of RNA to the protein. The turbidity assays showed that that IGF2BP1 S181E mutant formed condensates with slower kinetics indicating that phase separation is impaired for this mutant (**Fig. 2J**, **Table 4**, wild-type: t_1/2_ =167 s, S181E: t_1/2_ = 252 s). Moreover, the condensates formed by the IGF2BP1 S181E mutant showed lower scattering intensity compared to the wild-type protein (**Fig. 2J**, **Table 4**, wild-type: OD_480_ = 0.068, S181E: OD_480_ = 0.035), consistent with the microscopy data that showed formation of smaller condensates (**Fig. 2G, I**). In contrast, the Y396E mutant had slightly faster formation kinetics compared to the wild-type IGF2BP1 with similar scattering intensity (**Fig. 2J**, **Table 4**, Y396E: t_1/2_ =115 s, OD_480_ = 0,064,) Altogether, our data indicate that phosphomimetic mutations in the IGF2BP1 disordered linker regions impact the formation of RNP condensates in opposing directions and a context-dependent manner.

Condensate fluidity impacts fusion and growth and as a result affects condensate size ^45^. We wondered whether differences in the size of IGF2BP1 condensates formed by the phosphomimetic mutants stem from differences in their fluidity. Fluorescence recovery after photobleaching (FRAP) experiments measure protein diffusion coefficients and are widely used to measure condensate fluidity ^46, 47^. We tested whether IGF2BP1 phosphomimetic mutants had different diffusion dynamics in RNP condensates by FRAP experiments. We formed IGF2BP1-RNA condensates with sub-stochiometric mCherry-tagged IGF2BP1, and monitored recovery of mCherry fluorescence in the condensates after photobleaching. To be able to decouple condensate growth from fluorescence recovery and to allow formation of large condensates that are tractable for the FRAP measurements, we incubated IGF2BP1 with RNAs for 90 minutes before performing the photobleaching experiments. The FRAP data revealed that mCherry-IGF2BP1 fluorescence did not recover even 15 minutes after photobleaching, indicating that IGF2BP1 formed stable complexes with RNAs in the condensates. The long recovery time likely reflects the multivalent nature of IGF2BP1’s interaction with RNA which results in high residence times and long-lived interactions ^48^. Consistent with this explanation, FRAP experiments showed that similar to wild-type IGF2BP1, the condensates formed by IGF2BP1 phosphomimetic mutants did not recover fluorescence intensity even after 15 minutes (**Fig. 3A**).

**Figure 3.**
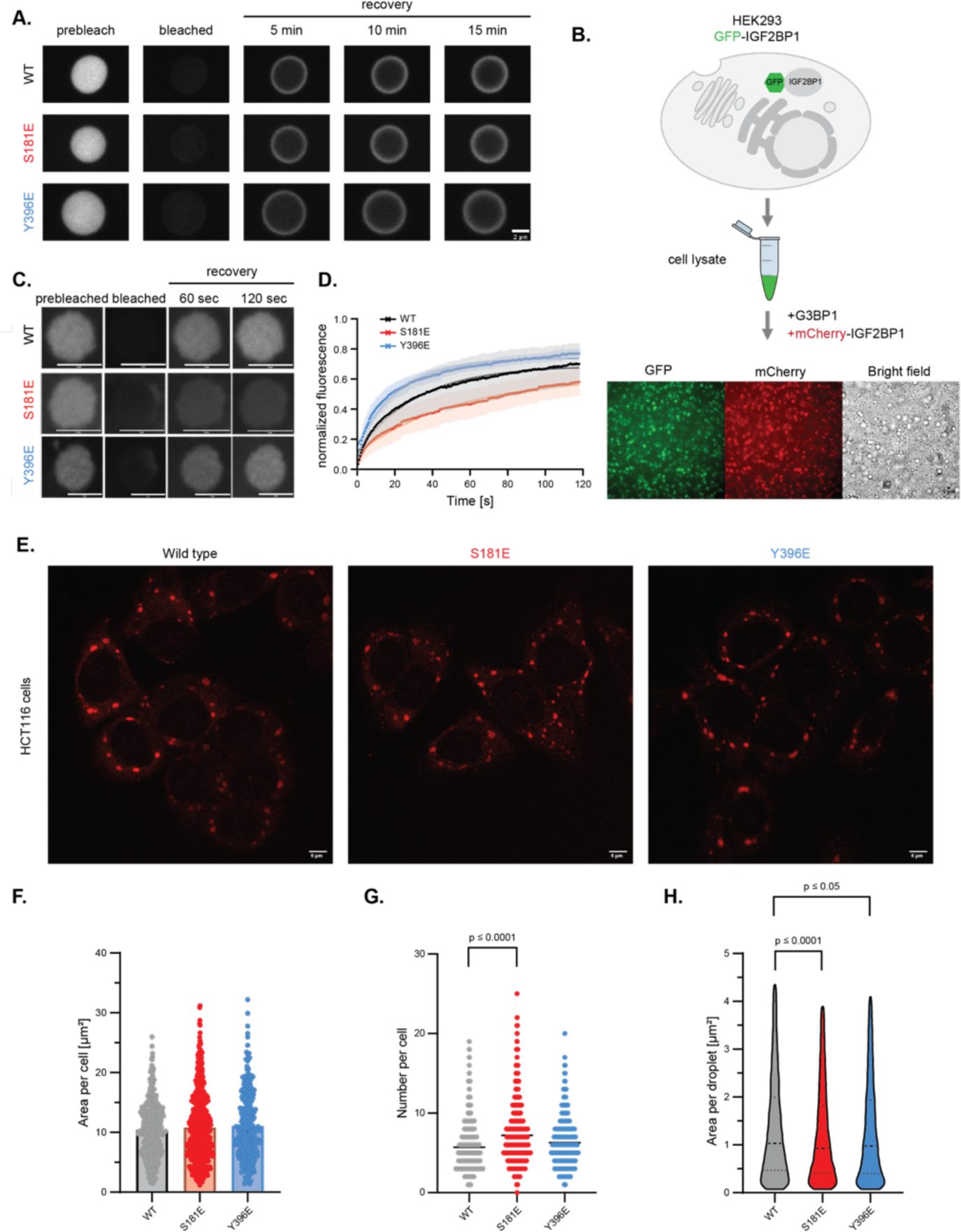
**(A)** Representative images of Fluorescence Recovery After Photobleaching (FRAP) of condensates formed by 5 µM full-length IGF2BP1 wild-type, S181E and Y396E and 5% mCherry-labeled IGF2BP1 constructs in the presence of 5 µM XBP1 36 nt RNA after 90 min incubation. Scale bar is 2 µm. **(B)** Schematic depiction of G3BP1-induced RNP granule formation in lysates and representative images of the incorporation IGF2BP1 into RNP granules. **(C)** Representative images of FRAP experiments with mCherry-IGF2BP1 wild-type, S181E and Y396E mutants in RNP granules after 100 min incubation. Scale bar is 5 µm. **(D)** Recovery curves of FRAP experiments with mCherry-IGF2BP1 wild-type (black), S181E (red) and Y396E (blue) mutants in G3BP1-induced granules after 100 min incubation. Error margins represent standard deviation. Fitted curve represents a one-phase association equation. n = 10 for wild-type, n=11 for S181E, n=13 for Y396E. **(E)** Representative fluorescence images of fixed HCT116 cells expressing mCherry-lIGF2BP1 wild-type, S181E or Y396E. Cells were fixed 60 min after stress induction by 500 µM arsenite. Scale bar is 5 µm. Quantification of condensates in HCT116 cells represented as scatter plots: **(F)** total area of condensates per single cell (n = 449 for wild-type, n= 741 for S181E, n=278 for Y396E, bar represents mean value) **(G)** number of condensates per single cell (bar represents mean value) **(H)** area per condensate (bars represent the median and 25 % and 75 % quartiles)

We hypothesized that due to competition with other RNAs and RBPs, IGF2BP1 might have more dynamic interactions with RNAs in a cellular environment. Therefore, we next reconstituted IGF2BP1-RNA granules under conditions that mimic the nature of RNP interactions in a complex environment. It has been recently shown that supplementing mammalian cell lysates with G3BP1, the RBP that drives stress granule assembly, results in the formation of RNP condensates which closely resemble stress granules in terms of protein and RNA composition ^49^. IGF2BP1 is a component of the stress granules and we used this method to reconstitute IGF2BP1-containing RNP condensates in cell lysates obtained from HEK293 cells expressing GFP-tagged IGF2BP1. Addition of recombinant G3BP1 to cell lysates induced formation of condensates that were positive for GFP fluorescence, indicating that recombinant G3BP1 leads to the formation of IGF2BP1-containing RNP granules in cell lysates (**Fig. 3B**). We confirmed these results by adding recombinant mCherry-tagged IGF2BP1 after the formation of stress granules, and found that the mCherry-tagged IGF2BP1 was sequestered into G3BP1 induced-RNP condensates (**Fig. 3B**). We used this experimental set up to assess the dynamics of wild-type mCherry-IGF2BP1 and its phosphomimetic mutants S181E and Y396E in RNPs by FRAP experiments. We found that wild-type IGF2BP1 formed condensates with a mobile fraction of 68.8% after RNP condensate formation for 100 minutes with a recovery half-time of 21.6 sec (**Fig. 3C,D**, **Table 5)**. Intriguingly, FRAP experiments performed with the IGF2BP1 S181E mutant showed a more than two-fold increase in recovery half-time (37.5 s), and a slight decrease in the mobile fraction (62.2 %) compared to the wild-type IGF2BP1. In contrast, the IGF2BP1 Y396E mutant showed a faster recovery (15.5 s) and a slightly higher mobile fraction (74.1 %) compared to the wild-type. These data revealed that while the S181E mutant forms more stable interactions in the RNP condensates, the Y396E forms more dynamic ones.

### IGF2BP1 phosphomimetic mutants impact size and number of IGF2BP1 RNP granules in cells

In order to study the impact of IGF2BP1 phosphorylation on its assembly into RNPs in cells, we established mammalian cell lines stably expressing mCherry-tagged wild-type human IGF2BP1 or its phosphomimetic mutants using a lentiviral transduction approach. For these experiments, we selected two mammalian cell lines, U2OS cells (human osteosarcoma) and HCT116 cells (colon carcinoma cells). HCT116 cells do not express IGF2BP1. To exclude the possibility of functional redundancy between IGF2BP paralogs, we knocked out IGF2BP2/3 in HCT116 cells using CRISPR-cas9 gene editing (**Fig. Supp 5A**). As IGF2BP1 expression levels might impact condensate formation, we used fluorescence-activated cell sorting (FACS) to select cells which expressed mCherry-tagged wild-type IGF2BP1 and its phosphomimetic mutants at similar levels. For the HC116 cells, we used FACS to select single clones to ensure similar expression levels between wild-type and the mutants (**Fig. Supp 5B, C**). In addition, we studied U2OS cells due to their extended morphology which is well-suited for microscopy experiments. To simultaneously monitor stress granules and IGF2BP1 condensates, we used engineered U2OS cells expressing GFP-tagged stress granule marker G3BP1 ^50^. We picked a population of U2OS cells where mCherry-IGF2BP1 and its mutants were expressed similar to the wild-type protein levels (**Fig. Supp 5D, E**).

We assessed whether, similar to the *in vitro* results, IGF2BP1 phosphomimetic mutants impact RNP condensate formation in cells by studying IGF2BP1’s assembly into stress granules. Treatment of cells with sodium arsenite resulted in stress granule formation, evidenced IGF2BP1 sequestration into by G3BP1 positive RNP granules (**Fig. Supp. 5F**). Quantification of the number, size and total area of mCherry-IGF2BP1 positive granules revealed that HCT116 cells expressing wild-type and phosphomimetic mutants showed similar total granule area per cell (**Fig. 3E, F**, **Table 6**). Notably, the S181E mutant formed a higher number of condensates per cell (**Fig. 3G**, **Table 6**, mean condensate number, wild-type: 5.7, S181E: 7.2) with a slightly smaller area per condensate (**Fig. 3H**, **Table 6**, median condensate area, wild-type: 1.1 µm², S181: 1.0 µm²) indicating that similar to what we found *in vitro*, the S181E forms smaller condensates in cells (**Fig. 3H**, **Table 6**). These results were similar in both HCT116 and U2OS cell lines, with the differences being more pronounced in the HCT116 cell line. U2OS cells expressing the S181E mutant showed a similar total area of condensates and higher number of condensates (**Fig. Supp. 5G-J**, **Table 6**, median total area per cell, wild-type: 10.3 µm², S181E: 10.3 µm², mean number of condensates per cell, wild-type: 18.5, S181E: 20.7). The IGF2BP1 Y396E mutant did not display large differences in size and but higher number of condensates in HCT116 cells (**Fig. 3E-H**, **Table 6**, median condensate area, Y396E: 1.1 µm², mean condensate number per cell, Y396E: 6.3). Instead, condensates formed by the Y396E mutant in U2OS cells were smaller compared to those formed by wild-type IGF2BP1 (**Fig. Supp. 5G-J, Table 6**, median condensate area, wild-type = 1.7 µm², Y396E =1.4 µm²). We speculate that the expression of the wild-type IGF2BP1 and other IGF2BP paralogs, as well as cell-type dependent differences in signaling cascades, could result in the heterogeneity observed in U2OS cells. Overall, we found that the S181E mutation impacts the formation of IGF2BP1-containing granules in cells.

### IGF2BP1 forms a compact conformation in solution

We aimed to elucidate how phosphorylation of IGF2BP1 at the disordered linkers mechanistically impacts its condensation into RNP granules. We hypothesized that linkers might form contacts with each other or with the folded domains contributing to multivalency in condensates, and that phosphorylation could affect these interactions. Thus, we monitored the conformational status of IGF2BP1 by Small Angle X-ray Scattering (SAXS) analyses. SAXS is a solution scattering method that provides low resolution structural information on overall shape of molecules. Analysis of SAXS scattering curves (**Fig. Supp. 6A**) showed that IGF2BP1 displays a maximal extension (D_max_) of around 20 nm and a radius of gyration (R_g_) of 4.18 nm. For flexible molecules such as IGF2BP1, scattering curves represent an average of conformational states sampled by the protein. The Kratky plot of IGF2BP1 highlighted the presence of flexibility in the protein as the curve does not converge to the s axis (**Fig. Supp. 6B**). This is a consequence of a lack of structure in the linker regions. We further analyzed SAXS data using the ensemble optimization method (EOM) to extract information of those states ^51^. EOM generates a random pool of structures based on the available structural data and amino acid sequence of the protein. EOM then selects the ensemble of structures that fits the experimental SAXS data best. To generate models for EOM for IGF2BP1, we provided high-resolution structures for the individual domains with the flexible linkers. The comparison of the radius of gyration (R_G_) and maximal distance (D_max_) distributions of the random pool of structures compared to the selected ensembles revealed that the selected ensemble displayed more compact structures compared to the random pool (**Fig. 4A, B**). These analyses suggest that IGF2BP1 is in a conformational equilibrium between extended and compact states in solution with higher a number of molecules found in the compact state at any given time. These data also indicate the presence of low affinity intramolecular contacts within the molecule, leading to its compaction.

**Figure 4.**
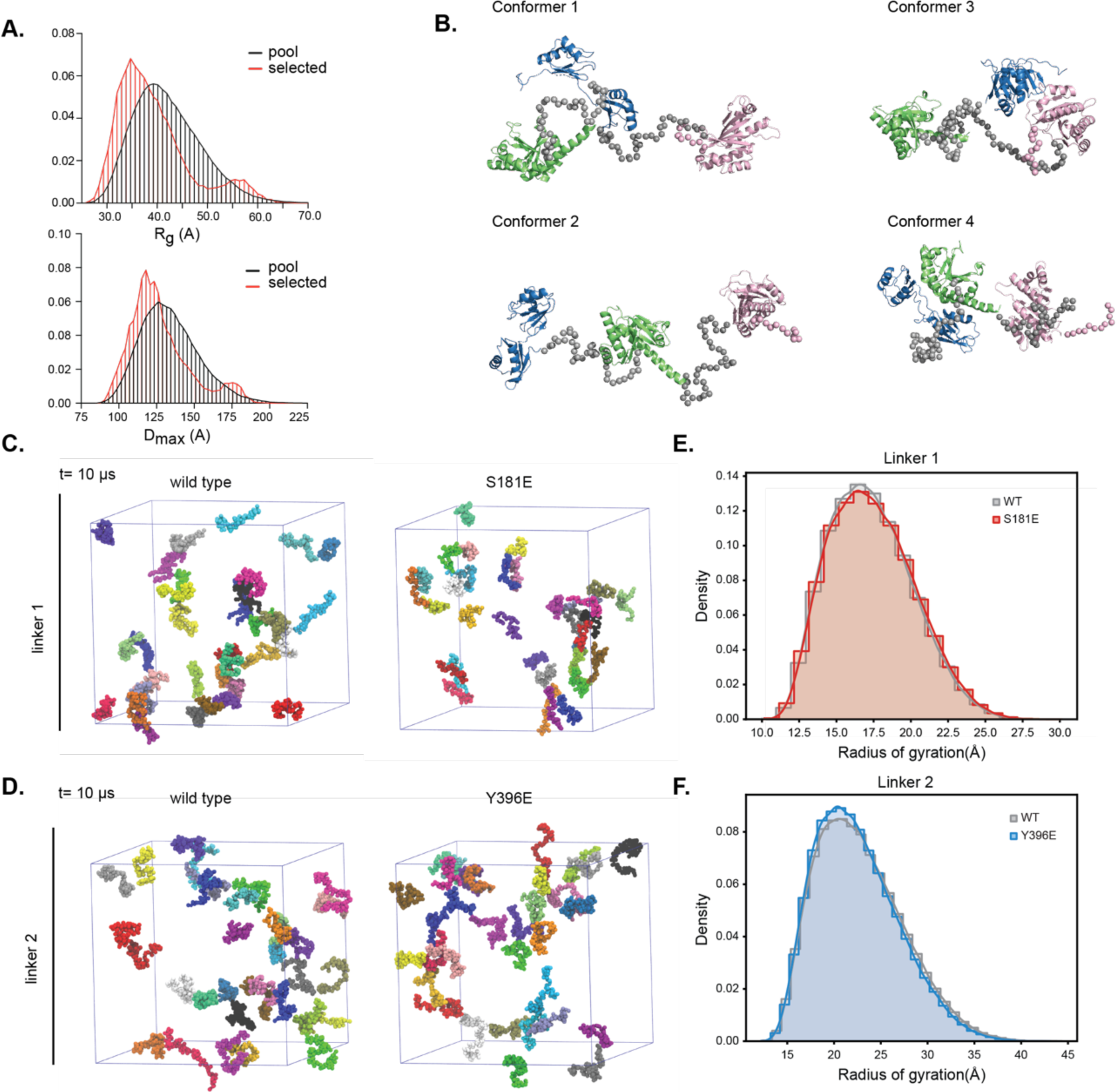
**(A)** Comparison of radius of gyration (R_g_, top) and maximum extension (D_max,_ bottom_)_ distribution of random conformations of IGF2BP1 (black) and selected pool (red) that best fit the experimental SAXS data based on EOM analyses. **(B)** IGF2BP1 structural conformers that best fit the experimental SAXS data (linkers are shown as gray spheres). The RRM1-2, KH1-2 and KH3-4 domains are colored in blue, green and pink, respectively. **(C)** Snapshot of coarse-grained Martini 3 Molecular Dynamics simulations of linker 1 and its mutant S181E at 10 µs. Each simulation box contains 33 copies of the polypeptide, depicted in different colors and sphere representation. Water and ions are not shown for clarity. (**D**) Snapshot of coarse-grained Martini 3 Molecular Dynamics simulations of linker 2 and its phosphomimetic mutant Y396E at 10 µs. Each simulation box contains 33 copies of the polypeptide, depicted in different colors and sphere representation. Water and ions are not shown for clarity. **(E)** R_g_ probability density distribution of linker 1 and its phosphomimetic mutant during MD simulations. **(F**) R_g_ probability density distribution of linker 2 and its phosphomimetic mutant during MD simulations.

Next, to assess whether phosphomimetic linker mutants impact the overall conformation of IGF2BP1 in solution, we used SAXS analyses. The SAXS data revealed that the scattering curves of the wild-type IGF2BP1 were very similar to the phosphomimetic mutants (**Fig. Supp. 6A**). In line with this data, EOM analyses of the phosphomimetic mutants showed comparable distributions of D_max_ and R_g_ values (**Fig. Supp. 6C, D**). The similarity of D_max_ and R_g_ values indicated that wild-type IGF2BP1 and phosphomimetic mutants share a comparable overall conformational ensemble. These data revealed that the negative charges introduced to the disordered linkers do not result in large conformational changes in the protein. As SAXS data provides low resolution information on a conformational ensemble, we next used orthogonal methods to study the impact of phosphorylation on the conformation and self-assembly of the linkers.

### IGF2BP1 linkers do not form intra and inter molecular contacts in isolation

Disordered regions contribute to phase separation of proteins. Therefore, the phosphorylation of the linkers could regulate condensate formation through modulating self-association between these regions. To monitor the condensation propensity of the linkers in isolation, we performed coarse-grained Molecular Dynamic (MD) simulations on the disordered linkers and their corresponding phosphomimetic mutants (**Fig. 4C-D**). We conducted Martini 3 simulations of multiple copies of the polypeptide chains of each linker under explicit solvent conditions and rescaled protein-water interactions in the force field to account for the disordered linkers. MD simulations revealed that over a 10 µsec time scale, none of the linkers assembled into clusters, suggesting that they do not display a strong propensity to self-associate. The phosphomimetic mutants behaved like their wild-type counterparts. These data are consistent with our *in vitro* experiments in which linkers did not form condensates at 150 µM even in the presence of a crowding agent (**Fig. Supp. 4M**).

To experimentally dissect this further, we performed Nuclear Magnetic Resonance spectroscopy (NMR) experiments. For NMR analyses, we produced ^15^N-isotope labeled proteins. ^1^H-^15^N Heteronuclear Single Quantum Coherence (HSQC) experiments revealed a small dispersion of backbone amide signals thus validating the disordered nature of the linker segments (**Fig. Supp. 7A-D**). We assessed self-association of the isolated disordered linkers by monitoring their NMR signal intensity at different linker concentrations. For monomeric non-self-associating molecules, NMR signal intensity should increase linearly with increases in protein concentration. In contrast, if the linkers associate with each another, this would result in broadening of the NMR signals and a drop in signal intensity over the concentrations where these interactions occur. We acquired HSQC spectra of the disordered linker 1 at protein concentrations ranging from 25-200 µM (**Fig. Supp. 7E**). Our data revealed that the signal intensity of the backbone amide groups increased linearly with increased protein concentration and there was no deviation from the predicted intensities. These data indicated that the disordered linker 1 does not self-associate under the conditions we tested. We made similar observations with the disordered linker 2 under the same experimental conditions (**Fig. Supp. 7F**). These data supported MD simulations showing that the intrinsically disordered linker regions do not self-associate in isolation up to 200 µM protein concentration.

### IGF2BP1 linkers adopt an extended conformation

The amino acid properties of interdomain linkers influence their effective solvation volume and impact the conformational space of linear multi-domain proteins ^52^. Importantly, linker compaction has been suggested to regulate phase separation properties in multi domain proteins^52^. As the linkers we tested did not have a propensity to self-associate, we next tested whether they form short distance intramolecular contacts that underlie the compaction we measured by SAXS analyses. These contacts might impact the phase separation propensity of IGF2BP1. We calculated the frequency of contacts formed within the same polymer chain (cis-interactions) in MD simulations. The contact maps obtained (**Fig. Supp. 6E, F**) indicated that cis-contacts are rarely established among residues separated by more than four amino acids within the disordered linker 1 and linker 2, and do not occur in the phosphomimetic mutants. These results highlight how cis-interactions are sparse in the wild-type as well as in the phosphomimetic mutants. We observed no long-range interaction within the chains. This was reflected in the lack of trans-interactions in our simulation box. In line with these observations, the distributions of the R_g_ of the linkers (linker 1 and linker 2) and the phosphomimetic mutants in MD simulations revealed that both linker 1 and linker 2 and their phosphomimetic mutants adopt expanded conformations with similar median R_g_ values (**Fig. 4E-F**). Importantly, both linker 1 and linker 2 display higher R_g_ values compared to an ideal polymer (Analytical Flory Random Coil) ^53^ with perfectly balanced polymer-solvent and polymer-polymer interactions and with the same amino acid sequence for each linker. These higher R_g_ distribution values correspond to a polymer model where polymer-solvent interactions are more prominent than polymer-polymer interactions (**Fig Supp. 7G, H**). Altogether, the MD simulations suggested that the linkers form an extended solvent-exposed conformation. Based on these data and the NMR experiments, we excluded the possibility that the differences in linker compaction lead to the different phase separation propensities observed for the IGF2BP1 phosphomimetic mutants.

### The linkers form low affinity interactions with the folded domains and RNA

Besides compaction and self-association of the linkers, transient interactions between the linkers and folded domains could lead to the IGF2BP1 compaction observed in SAXS experiments. To test whether the linkers are able to interact with the folded domains of the protein, we performed ^1^H-^15^N HSQC experiments for the titration series of the ^15^N-labeled linkers with the NMR invisible domains (RRM1-2, KH1-2 and KH3-4). Interactions between the linkers and domains are expected to induce chemical shifts perturbations (CSPs) and/or peak broadening which results in a reduced signal intensity for amino acids in close proximity to the interaction surface. To assign the signals in the NMR spectra of the linkers, we used three-dimensional sequential assignment strategy and we were able to unambiguously assign 70-80% of the signals in both linkers.

The titration of linker 1 with the folded domains showed very low CSPs (< 0.015 ppm) that were consistently alike for both the wild-type and S181E mutant protein (**Fig. 5A**, **Fig. Supp. 8A-I**). Therefore, we concluded that linker 1 does not interact with the folded domains of the protein. Besides forming protein-protein interactions, the disordered linkers could potentially form contacts with RNA which might modulate the phase separation propensity of IGF2BP1. Linker 1 contains an RGG/RG motif (aa 167-RRGGFGSRG-175, **Fig. 5A**), which has been shown to bind RNA non-specifically in other RBPs ^54^. Therefore, we investigated whether linker 1 could contribute to protein-RNA interactions by measuring HSQC spectra of linker 1 in the presence of a model RNA (12xUG) that is recognized by RGG containing proteins through formation of an RNA quadruplex^55^ and a 10 nt RNA derived from XBP1. Titration of the wild type linker 1 with the 12xUG RNA resulted in large chemical shift perturbations and decrease in signal intensity of the residues around the RGG-RG motif (G170, G172 and G175, **Fig. 5B-D** and **Fig. Supp. 8J, K**). Notably, the linker 1 phosphomimetic mutant S181E was impaired in its binding to RNA evident as lower CSPs and lower drop in the signal intensity around the phosphorylation site upon its titration with the 12xUG RNA. We observed similar, albeit weaker, CSPs of the same residues upon titration of the linker 1 with the 10nt XBP1 RNA indicating that this RNA bound to linker 1 with a much lower affinity (**Fig. Supp. 8L-P**). The difference between the wild-type and the S181E mutant was more pronounced due to low affinity interaction (**Fig. Supp.9L-P**). Our data suggest that impairing non-specific RNA binding by linker 1 via phosphorylation tunes the formation of IGF2BP1-containing RNP condensates.

**Figure 5.**
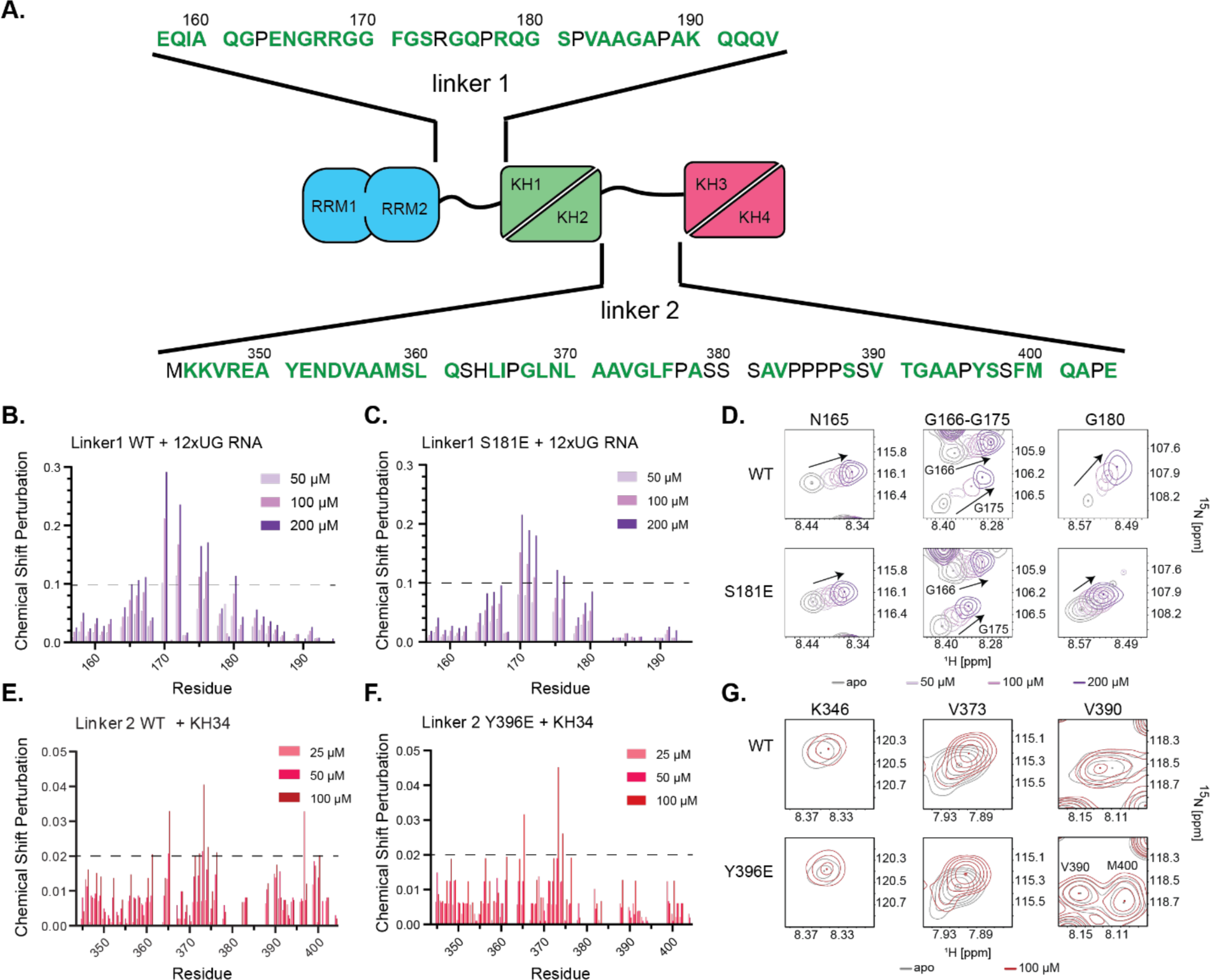
**(A)** Position and amino acid sequence of wild-type linker 1 and linker 2. **(B)** Chemical Shift perturbations (CSP) analyses of ^15^N-labeled wild-type linker 1 in the absence (black) and presence of various concentrations of 12xUG RNA (shades of purple) in ^15^N-^1^H HSQC experiments. **(C)** CSP analyses of ^15^N-labeled linker 1 S181E mutant in the absence presence of various concentrations of 12xUG RNA in HSQC experiments. **(D)** Representative backbone amide signals from wild-type linker 1 and the S181E mutant in absence and the presence of various concentrations of 12xUG RNA from HSQC spectra. **(E)** CSP analyses of ^15^N-labeled wild-type linker 2 in the absence and presence of different concentrations of KH3-4 in HSQC experiments. **(F)** CSP analyses of ^15^N-labeled linker 2 Y396E mutant in the absence and presence of various concentrations of KH3-4 in HSQC experiments. **(G)** Representative signals of ^15^N-labeled wild-type linker 2 and Y396E in the absence and presence of 100 µM KH3-4.

We next assessed whether linker 2 interacts with the folded domains of IGF2BP1 by NMR experiments. The NMR analyses revealed that titration of wild-type linker 2 with RRM1-2 as well as the KH1-2 domains showed small CSPs around aa S388-P395 (aa 388-SSVTGAP-395) (CSP > 0.015) (**Fig. Supp. 9A-F**) indicating that linker 2 interacts with these domains with a low affinity in the millimolar range. Importantly, titration of the linker 2 phosphomimetic Y396E mutant with both domains displayed reduced CSPs in this region. These data demonstrate that the Y396E mutation modulates low affinity binding of linker 2 to RRM1-2 and KH1-2 domains.

Titration of linker 2 with KH3-4 domains displayed the largest CSPs observed for all folded domains, revealing that linker 2 most strongly interacts with KH3-4 dimers (**Fig. 5E-G, Fig. Supp. 9G-H**). Nevertheless, the fluorescence anisotropy experiments showed that a construct containing both linker 2 and the KH3-4 domains (linker 2-KH3-4) bound to RNA with a similar affinity to the KH3-4 domains alone. This result indicates that the low affinity interaction between linker 2 and the KH3-4 domains does not impair RNA binding (**Fig. Supp. 9I**). The KH3-4 pseudodimers bound to a largely hydrophobic region in linker 2 covering Q361-F376 (aa 361-QSHLIPGLNLAAVGLF-376). Notably, of all IGF2BP1 domains this segment only bound to KH3-4, emphasizing the specificity of this interaction. In addition, KH3-4 domains bound to a region covering the Y396 phosphorylation site (aa V390-M400). The linker 2 Y396E phosphomimetic mutant showed reduced CSPs upon KH3-4 binding close to the mutation site, yet the interaction of KH3-4 with the hydrophobic segment (aa Q361-F376) was not affected by the Y396E mutation. To sum up, we found that linker 2 forms low affinity contacts with KH3-4 pseudodimers, which are modulated in the phosphomimetic mutant.

We anticipate that in full length IGF2BP1, linker 2 binds more tightly to the KH3-4 pseudodimers as forced proximity in the polypeptide chain increases their local concentration. To test this possibility, we compared the NMR spectrum of linker 2 alone with that of KH1-4 and L2-KH3-4 constructs under experimental conditions that only detect disordered regions due to their high dynamics and signal intensity (**Fig. Supp. 9J**). The NMR signals from the disordered regions in KH1-4 largely overlapped with that of the L2-KH3-4 construct, suggesting that presence of KH1-2 does not change conformation of linker 2. In contrast, the NMR signals of linker 2 alone only partially overlapped with both of the spectra. The disordered regions in L2-KH3-4 construct mainly consists of the linker 2, therefore these results indicated that the linker 2 forms contacts with KH3-4 pseudodimers in cis, supporting our model.

Overall, the NMR data revealed that while linker 1 binds to RNA, linker 2 forms low affinity interactions with the folded domains in IGF2BP1 and both of those interactions are modulated in the relevant phosphomimetic mutants. We anticipate that these low affinity interactions are more pronounced in the condensed phase with high protein and RNA concentration (up to 20 mM)^56^, resulting in the prominent effect of these mutants in IGF2BP1 RNP condensates. Therefore, our data converge on the model that the proteotoxic stress-regulated phosphorylation of S181 increases the rigidity and decreases the size of IGF2BP1 containing RNP granules by abrogating low-affinity protein-RNA interactions while Y396E phosphorylation increases the dynamics and size of these condensates by impairing self-association and protein-protein interactions (**Fig. 6**).

**Figure 6.**
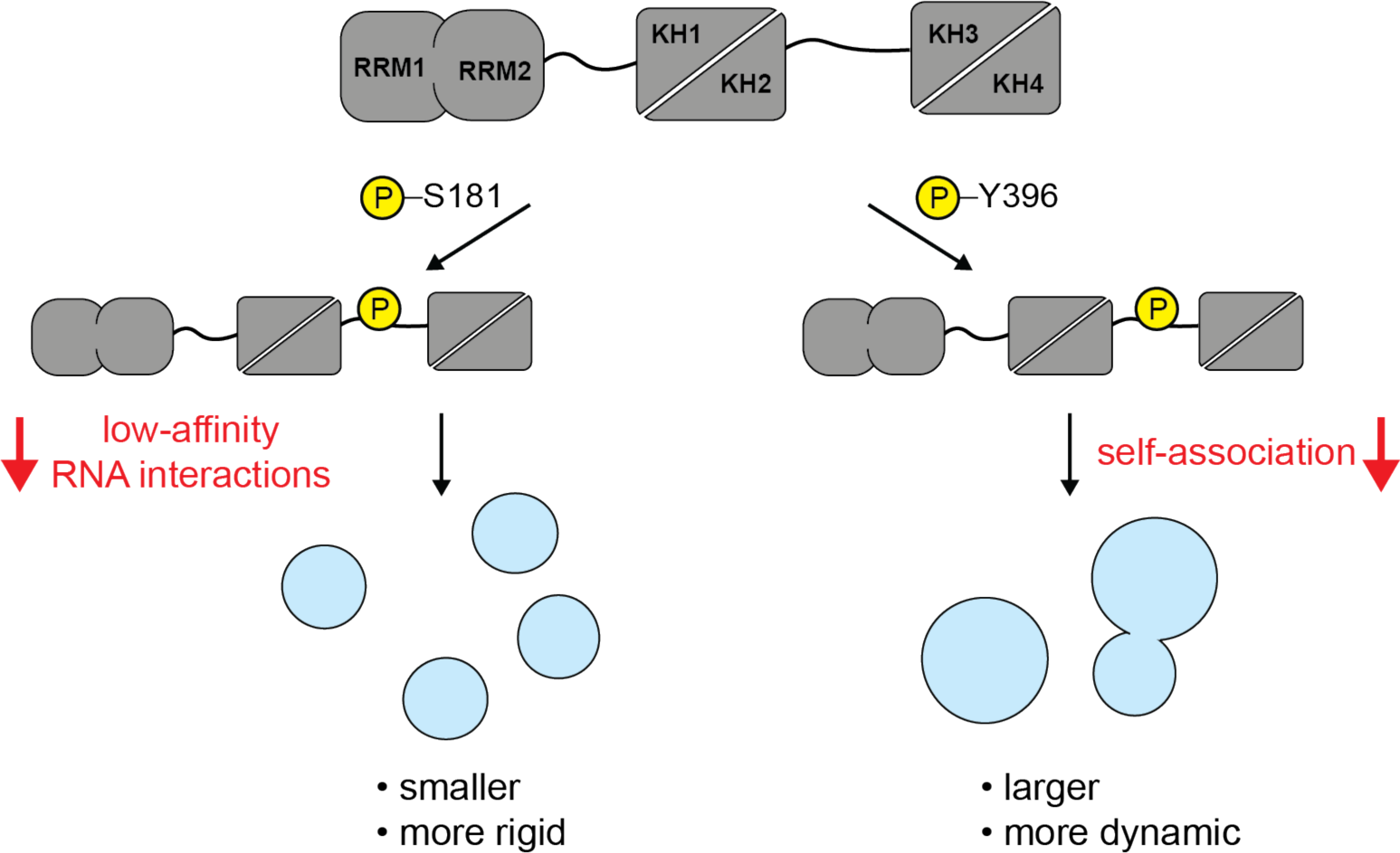
Model of the effect of linker 1 and linker 2 phosphorylation on IGF2BP1 condensate formation.

## Discussion

Posttranslational modifications (PTMs) regulate protein function in a reversible, tuneable manner that allows exquisite spatiotemporal control ^57^. PTMs regulate many RBPs by modulating target RNA binding, interaction with partners, or subcellular localization and thereby contribute to control RNA metabolism in cells ^58–60^. It has become increasingly clear that PTMs regulate assembly of RBPs into biomolecular condensates by phase-separation. Here, using mass spectrometry, we mapped steady-state and stress-induced phosphorylation sites in IGF2BP1.

Targeted proteomics identified novel stress-dependent phosphorylation sites in IGF2BP1. Apart from the S181 site, whose phosphorylation decreased approximately two-fold during ER stress, ER stress only mildly impacted the phosphorylation status of IGF2BP1. Notably, the oxidative stress resulted in an increase in phosphorylation at various sites throughout the protein including a in a two-fold increase in S181 phosphorylation revealing context dependent nature of this phosphorylation event. IGF2BP1 residue S181 was previously proposed to be phosphorylated by mTORC2 ^31, 32^. Importantly, recent data also predicted that the S181 site is phosphorylated by the CMKG kinase family ^37^. Several members of the CMGC kinase family are activated during oxidative stress and the family member DYRK3 partitions to stress granules and regulates their disassembly ^61^. Under the induced ER stress conditions used here, cells did not form stress granules, suggesting that their formation might be important for stress-induced IGF2BP1 phosphorylation at this site. Apart from prominent S181 phosphorylation (64% of the total protein pool), most of the identified phosphorylation sites in IGF2BP1 were only modified at sub-stoichiometric levels (< 1%). We anticipate that those sites might be regulated in a spatial-or cell-type-specific manner and could be present at a higher frequency in other cell types.

In the absence of RNA, purified IGF2BP1 displayed monodisperse, monomeric behavior. In contrast, in the presence of RNAs with multiple IGF2BP1 binding motifs, IGF2BP1 assembled into RNP condensates. By systematic analyses of IGF2BP1 truncation mutants, we revealed that the KH3-4 domains in IGF2BP1 drive the formation of RNP condensates. These data are in line with experiments in which only RNA-binding mutants of KH3-4 domains impaired IGF2BP assembly into stress granules in cells ^39^. Minimal IGF2BP1 RNP condensates consisting of RNA and IGF2BP1 were highly rigid due to multivalent interaction of the RNA with the multidomain protein IGF2BP1. Notably, while full length IGF2BP1 formed droplets, KH1-4 domains formed meshed networks, underlining the role of RRM domains in increasing dynamics within IGF2BP1-RNA assemblies. These data suggest that the competing non-specific interactions of the highly promiscuous RRM1-2 pseudodimers increase the dynamics in RNA-protein interactions in the IGF2BP1 condensates. In line with these observations, IGF2BP1 RNP condensates examined *in vitro* in cell lysates displayed increased dynamics. We anticipate that in cells other RBPs compete with the available RNA-binding sites and this way weaken the interactions between RBPs and RNAs increasing the dynamics of RNA-protein interactions within condensates.

Systematic biochemical analyses with truncation and point-mutants revealed that the IGF2BP1 KH3-4 domains bind to RNA with the highest affinity, with KH3 showing the strongest contribution. In contrast to what was shown for the IGF2BP1 chicken homologue ZBP1, we did not observe the looping of RNA around the KH3-4 domains which results in an avidity effect for both the *ACTB*- and *XBP1*-derived RNAs ^41, 42^. We instead observed an avidity effect driven by combined binding of the KH3-4 and KH1-2 domains to RNA. Importantly, RNA-binding mutants of both the KH3 and KH4 domains in full length IGF2BP1 still bound tightly to RNA through avidity driven by RRM1-2 and KH1-2 domains. These data support earlier findings for the IGF2BP3 paralog underlining the importance of combinatorial recognition of the IGF2BP targets ^20, 21^. We propose that the KH3-4 domains dock onto the RNA with medium affinity and fast kinetics and an avidity effect driven by the KH1-2 and RRM1-2 domains increases the affinity and specificity of IGF2BP1 binding to its targets. Phosphomimetic mutants of IGF2BP1 linkers bound to short model RNAs containing two IGF2BP1 binding motifs with an affinity similar to the wild-type protein. Notably, phosphomimetic mutants showed slight differences in their binding to the 200 nt-long *EIF2A*- and *XBP1*-derived RNAs in EMSA assays. These data suggested the possibility of RNA-dependent differences for IGF2BP1 phosphorylation. Altogether, our data converge on the model that linker phosphorylation does not abolish IGF2BP1’s interaction with RNAs, yet might modulate IGF2BP1-RNA assemblies in cells as proposed earlier ^7, 32^.

Combining cell biology and *in vitro* reconstitution methods, we found that IGF2BP1 phosphorylation at the disordered linkers (S181, Y396) modulates its assembly into RNP granules. Intriguingly, while phosphorylation of S181 at linker 1 decreases the dynamics and the size of IGF2BP1 RNP condensates, phosphorylation of Y396 showed an opposite effect *in vitro* highlighting the regulatory potential of these sites. Using structural methods, we dissected how phosphorylation of the linkers modulates the conformational state of IGF2BP1 and thereby formation of RNP condensates. We hypothesized that linkers might be regulating IGF2BP1 condensation by three possible mechanisms; (i) self-association (ii) interaction with folded domains or (iii) interaction with RNAs. The NMR experiments, *in vitro* reconstitution and MD simulations showed that linker 1 and linker 2 do not display a propensity to self-associate. Instead, NMR experiments revealed that while linker 1 does not interact with folded domains of the IGF2BP1, it formed low affinity contacts with RNA through its RGG/RG motif. We propose that these low affinity contacts increase the dynamics of IGF2BP1 in the RNP condensates. Importantly, these low affinity interactions were impaired in the phosphomimetic mutant S181E. FRAP experiments showed that IGF2BP1 S181E does not recover as fast as the wild-type protein indicating that it forms more stable assemblies in the condensates in comparison to wild-type IGF2BP1. Our results converge on the model that IGF2BP1 S181 phosphorylation impairs low affinity non-specific interactions of the linker with the RNA. Lack of competing low affinity interactions results in more stable binding of S181E with RNA in the condensates that is observed as decreased fluidity in FRAP experiments. The hardening of condensates impairs condensate fusion and leads to formation of smaller condensates which we observed *in vitro* and in cells for the S181E mutant. We anticipate that hardening of the condensates through S181 phosphorylation might serve functional roles in cells by preventing partitioning of certain factors into the RNP granules to regulate RNA metabolism. Alternatively, hardening of condensates may cage RNAs preventing their translation or might provide structural rigidity that ensures integrity during transport of granules. Indeed, the non-phosphorylatable mutant of IGF2BP1 S181A is impaired in its capacity to stabilize c-Myc RNA in cells^34^ and impaired its dendritic distribution and motility^33^. In addition to phosphorylation, RGG sequences are methylated by protein arginine methyltransferase 1 (PRMT1), which can impact RNA binding of linker 1 ^62^. Therefore, the linkers might be subjected to more intricate regulation in cells in a context and tissue-specific manner.

Using NMR spectroscopy, we found that unlike linker 1, linker 2 forms low affinity contacts with all folded domains in IGF2BP1, showing the strongest binding to the KH3-4 dimers. Notably, all folded domains bound weakly to the C-terminal segment (aa V390-M400) in linker 2 covering the Y396 phosphorylation site and those interactions were impaired for the Y396E phosphomimetic mutant. Those data uncovered that the molecular contacts between linker 2 and the folded domains were partially impaired in the Y396E mutant. FRAP experiments showed that the IGF2BP1 Y396E mutant is more dynamic in condensates compared to the wild-type protein. Our data suggest that changes in low affinity interaction networks formed between linker 2 and folded domains in the Y396E mutant slightly increases IGF2BP1 dynamics in RNP condensates. We propose that increased dynamics facilitate the formation of condensates which we observed as larger condensates *in vitro*. Notably, the IGF2BP1 Y396E mutant expressed in cells did not show consistent effects on the size of IGF2BP1 granules, most likely due to high heterogeneity of the stress response pathways or increased compositional heterogeneity of the RNPs in cells compared to the *in vitro* experiments.

Our findings highlight a highly tunable regulatory mechanism where modulation of low affinity interactions through phosphorylation in turn impacts the physical properties of RBPs in RNP condensates. Our data exemplifies how the effects of PTMs are amplified in the condensate environment, thus providing an increased regulatory capacity to control biomolecular interactions in membraneless organelles.

## Acknowledgements

We are grateful to the late Thomas Peterbauer at the Max Perutz Labs Biooptics Light Microscopy Facility for his help and support. We thank Mila Asparuhova, Gizem Celebi and Isabell Niedermoser for their technical support and help. We appreciate the support from Julia Scholz in image analysis. We thank Gijs Versteeg and his lab for their help with lentiviral transduction and providing us with the expression plasmids. We thank Kitti Csalyi and Thomas Sauer at Max Perutz Labs Biooptics FACS facility for their help. We thank Georg Kontaxis for his continuing support with NMR measurements. Proteomics analyses were performed by the Mass Spectrometry Facility at Max Perutz Labs using the VBCF instrument pool. We are grateful to Max Perutz Labs Mass spectrometry facility, Markus Hartl, Dorothea Anrather, Wei-qiang Chen and David Hollenstein for their help and support with measurements, data analyses and experimental design. We are grateful to Thomas Leonard for his help with SEC-SAXS measurements and his invaluable input in data analyses. We thank Jeffrey A. Chao for kind gift of MBP-tagged IGF2BP1 constructs. We acknowledge funding from Austrian Science Fund (FWF-SFB F79 and FWF-W 1261) to GEK. E.S. and R.C. acknowledge support and funding by the Frankfurt Institute of Advanced Studies, the LOEWE Center for Multiscale Modelling in Life Sciences of the state of Hesse, the Collaborative Research Center 1507 “Membrane-associated Protein Assemblies, Machineries, and Supercomplexes” (Project-ID Project ID 450648163 – P09), and the International Max Planck Research School on Cellular Biophysics (to R.C.), the Centre for Scientific Computing of the Goethe University and the Jülich Supercomputing Centre for computational resources and support. The research of T.M. was supported by Austrian Science Fund (FWF) grants P28854, I3792, DOC-130, and DK-MCD W1226; Austrian Research Promotion Agency (FFG) grants 864690 and 870454; the Integrative Metabolism Research Center Graz; the Austrian Infrastructure Program 2016/2017; the Styrian Government (Zukunftsfonds, doc.fund program); the City of Graz; and BioTechMed-Graz (flagship project). The GFP-G3BP1 engineered cell lines are kind gift of Witold Szaflarski, Pavel Ivanov and Paul Anderson.

## Materials and Methods

### *In vitro* construct cloning

All mCherry-IGF2BP1 constructs were cloned into a pET-47 plasmid with an N-terminal mCherry using Gibson assembly. Single point mutations were generated through site-directed mutagenesis as well as Gibson assembly. IGF2BP1 (Uniprot: Q9NZI8) mutation involved changing the wild-type Ser181 or Tyr396 residues to glutamic acid. In order to establish U2OS and HCT116 cell lines stably expressing IGF2BP1, a vector that contains hPGK promoter was used. The promoter, together with NheI restriction site was introduced into the pLX303 expression vector using XheI and BsrgI restriction enzymes. mCherry-tagged mutants of IGF2BP1 were amplified with pLX303_IGF2BP1_R (5’-CTCGCTAGCTCACTTCCTCCGTGCC-‘3) and pLX303_mCherry_F (5’-CTCACCGGTGCCACCATGGTGAGCAAGG-‘3) primers and incorporated into the modified pLX303 using the AgeI and NheI restriction sites. Sequencing confirmed IGF2BP1 integration.

### Cell culture

U2OS cell lines used in this study were grown in DMEM high glucose medium (Sigma) with 10% Fetal Bovine Serum (Gibco), 2 mM Glutamine (Sigma), 1% Pen/Step (Sigma). HCT116 conditionally expressing Tet-OsTIR1 were obtained from Masato Kanemaki Lab (Natsume et al. 2016) and grown in McCoy’s 5A (Modified) medium supplemented as above. The cells were maintained at 37°C, with 5% CO2, and were used for all biochemical experiments, live cell imaging, and Immunofluorescence. The cells were tested for mycoplasma contamination which was not detected.

### Establishment of IGF2BP2 and IGF2BP3 knockout cell lines

For knockout cell line generation, gRNA sequences (IGF2BP2: 5’-GAGCTGCCGGAGGTCGTCGG-3’; IGF2BP3: 5’-ACGCGTAGCCAGTCTTCACC-3’) were cloned into the pSpCas9 (BB)-2A-GFP (PX458) (plasmid #48138; Addgene, (Ran et al. 2013)). Cells were transiently transfected using jetOPTIMUS reagent (Tamar, 101000051) and GFP-positive single-cell clones were FACS sorted at BD FACSAria IIIu at Max Perutz Labs BioOptics FACS Facility.

### Mass spectrometry– Sample Preparation

For mass spectrometry of IGF2BP1, we used HEK293T cells expressing IGF2BP1 tagged with split-GFP at the endogenous locus. These cells were the kind gift of Manuel Leonetti (Chan Zuckerberg BioHub, USA ^63^). Briefly, the cells were generated by integrating GFP^1–10^ into HEK293T cells by lentiviral integration. This cell line was then used to introduce GFP^11^ into the IGF2BP1 using CRISPR-Cas9 gene editing approaches ^64^. For the immunoprecipitation experiments, HEK293T cells were treated with the respective diluent as a control (DMSO for thapsigargin and tunicamycin or PBS for sodium arsenite). To induce stress, cells were treated with 250 µM sodium arsenite for 1 and 2 hrs, with 5 µg/mL tunicamycin for 4 hrs and with 250 nM thapsigargin for 1.5 and 4 hrs. IGF2BP1 was immunoprecipitated from HEK293T cells by GFP-trap magnetic beads (ChromoTek). The cells were lysed using cold lysis buffer (20 mM HEPES pH 7.4, 150 mM NaCl, 0.5 mM EDTA, 0.1% SDS, 0.5% Triton X-100, 0.2% Deoxycholate, 2x Complete™, EDTA-free Protease Inhibitor Cocktail (Roche), 0.5 mM PMSF and 1x PhosSTOP™ phosphatase inhibitor (Roche)). 25 µL of bead slurry was used for around 50 million HEK293T cells. The lysate was incubated with the GFP-trap beads for 2 hrs at 4°C to allow binding of the protein. After binding, the beads were washed 5 times with 1 mL of ice-cold wash buffer (20 mM HEPES, 500 mM NaCl, 0.5 mM EDTA, 0.1% SDS, 0.5% Triton X-100, 0.2% Deoxycholate, 1x Complete™, EDTA-free Protease Inhibitor Cocktail (Roche)). Protein was eluted in 50 µL of 1x SDS sample buffer (50 mM Tris-HCl pH 6.8, 2% SDS, 0.1% Bromophenol blue, 10% glycerol+ 20 mM DTT) at +70°C for 10 min. The eluate was loaded on SDS-PAGE and stained by Colloidal Coomassie ^65^. The band corresponding to IGF2BP1 was cut and subjected to mass spectrometry analyses at the Max Perutz Labs Mass Spectrometry Facility.

For mass spectrometry of IGF2BP3, HEK293T cells expressing IGF2BP3-tagged with split-GFP at the endogenous locus and HCT116 cells were used. IGF2BP3 from HEK293T split-GFP cell lines were immunoprecipitated under same experimental conditions as IGF2BP1 described above. For the immunoprecipitation experiments, both cell lines were treated with DMSO or PBS as control. To induce stress, cells were treated with 250 µM of sodium arsenite for 1 or 2 hours, or 250 nM of thapsigargin for 1.5 or 4 hours or 5 µg/ml tunicamycin for 4 hours. For mass spectrometry of IGF2BP3 from HCT116 cells, we used antibodies against endogenous IGF2BP3. For immunoprecipitation three 15 cm (diameter) dishes of 60% confluent HCT116 per condition (around 50 million cells) were washed in ice-cold PBS, scraped, pelleted, and resuspended in 750 μL of ice-cold lysis buffer (25 mM HEPES pH 7.3, 150 mM NaCl, 0.5% NP-40, 0.5 mM EDTA, 10% Glycerol, 0.1% SDS, 0.2% Sodium Deoxycholate, 2x Complete™, EDTA-free Protease Inhibitor Cocktail (Roche), and 1x PhosSTOP™ phosphatase inhibitor (Roche)). Cells were lysed by incubation with the lysis buffer on ice for 15 min with intermittent vortexing and passing the cell suspension three times through a 25G needle. The lysate was clarified using two-step centrifugation for 5 min at 1,000 g and for 15 min at 13,000 g, and treated with 1 U/μL RNase T1 (Thermo Scientific) rotating at room temperature for 15 min. For IP from three 15 cm (diameter) dish 30 μg of anti-IGF2BP3 antibody was coupled to Dynabeads as described before in 1 μg: 4 µL antibody: beads ratio. We used the MBL antibody (RN009P, lot 005) for the samples treated with tunicamycin and the respective 0.001% DMSO controls, and the Proteintech (14642-1-AP, lot 00090203) antibody for the thapsigargin, 0.0002% DMSO, arsenate and untreated control samples. The lysates were rotated at +4°C for 4 hours for the IP. The unbound fraction was removed using a magnetic rack and the immunoprecipitated complexes were washed five times in 1 mL of ice-cold high salt wash buffer (25 mM HEPES pH 7.3, 400 mM NaCl, 0.5% NP-40, 0.5 mM EDTA, 10% Glycerol, 1x protease inhibitors cocktail) with 3-min incubations on ice. Protein was eluted in 50 µL of 1x SDS sample buffer (50 mM Tris-HCl pH 6.8, 2% SDS, 0.1% Bromophenol blue, 10% glycerol) without DTT at +70°C for 10 min. DTT at 20 mM concentration was added to the collected eluates and the samples were heated at +70°C for 10 min. Samples were separated by SDS-PAGE and stained with colloidal Coomassie. The band corresponding to IGF2BP3 was cut from the gel and submitted for tandem mass spectrometry at the Max Perutz Labs Mass Spectrometry Facility.

### Mass Spectrometry – Sample processing

Coomassie stained gel bands were cut and destained with a mixture of acetonitrile (ACN) and 50 mM ammonium bicarbonate. Disulfide bridges were reduced using dithiothreitol and free SH-groups were subsequently alkylated by iodoacetamide. The digestion with trypsin was carried out overnight at 37 °C and was stopped by adding 10% formic acid to an end concentration of approximately 5%. Peptides were extracted from the gel with 5% formic acid by repeated sonication.

### Mass Spectrometry - Nano LC-MS/MS Analysis

Peptides were separated on an Ultimate 3000 RSLC nano-HPLC system using a pre-column for sample loading (Acclaim PepMap C18, 2 cm × 0.1 mm, 5 μm), and a C18 analytical column (Acclaim PepMap C18, 50 cm × 0.75 mm, 2 μm, all HPLC parts Thermo Fisher Scientific), applying a linear gradient from 2% to 35% solvent B (80% ACN, 0.08 % formic acid; solvent A 0.1 % formic acid) at a flow rate of 230 nL/min over 60 min. Eluting peptides were analyzed on a Q Exactive HF-X Orbitrap mass spectrometer coupled to the HPLC via Proxeon nano-spray-source (all Thermo Fisher Scientific) equipped with coated emitter tips (New Objective).

The mass spectrometer was operated in data-dependent acquisition mode. Survey scans were obtained in a mass range of 375-1500 m/z with lock mass on, at a resolution of 120000 at 200 m/z and a normalized AGC target of 3E6. The 10 most intense ions were selected with an isolation width of 1.6 m/z, for max. 200 ms at a normalized AGC target of 1E5, and then fragmented in the HCD cell at 28% normalised collision energy. Spectra were recorded at a resolution of 30000. Peptides with a charge of +1 or >+6 were excluded from fragmentation; the peptide match feature was set to “preferred” and the exclude isotope feature was enabled. Selected precursors were dynamically excluded from repeated sampling for 30 seconds.

For the parallel reaction monitoring (PRM) analysis survey scans were acquired in a mass range of 375-1500 m/z with lock mass off, at a resolution of 30000 at 200 m/z and a normalized AGC target of 3E6. The PRM parameters - precursor m/z and retention time - were built based on the gel samples measured with DDA. Precursors of 33 peptides of interest (14 phosphorylated peptides and their unmodified counterparts plus 5 reference peptides) were isolated in a 0.7 m/z window and fragmented with 28% HCD collision energy. Orbitrap resolution was set to 30000, the normalized AGC target to 2E5. Maximal injection time for modified peptides was set to 200 ms.

### MS data analysis for identification of phosphorylation sites

The RAW MS data were analyzed with FragPipe (20.0), using MSFragger (3.8) ^66^, IonQuant (1.9.8) ^67^, and Philosopher (5.0.0)^68^. The default FragPipe workflow for label free quantification (LFQ-MBR) was used, except that “Add MaxLFQ”, “Match between runs (MBR), and “Normalize intensity across runs” were turned off. Cleavage specificity was set to Trypsin/P, with two missed cleavages allowed. The protein FDR was set to 1%. A mass of 57.02146 (carbamidomethyl) was used as fixed cysteine modification; methionine oxidation, protein N-terminal acetylation, and serine/threonine/tyrosine phosphorylation were specified as variable modifications. MS2 spectra were searched against the *H. sapiens* reference proteome from Uniprot (Proteome ID: UP000005640, release 2023.03) containing 20598 entries, concatenated with a database of 379 common laboratory contaminants (in house database).

Computational analysis was conducted using Python along with two in-house Python libraries, “MsReport” (version 0.0.20) and “XlsxReport” (version 0.0.6) ^69^. To compile a list of confidently identified phosphorylation sites for IGF2BP1, IGF2BP2, and IGF2BP3, we utilized the individual “ion.tsv” tables generated by FragPipe. Initially, the ion tables were concatenated and non-phosphorylated peptide ions were filtered out. Subsequently, multiple phosphorylated peptide ions were selected, and separate entries were generated for each phosphorylated site by duplication. The specific site localization probability was then extracted for each of the duplicated site entry. Entries with a peptide probability lower than 95% or a phosphorylation site localization probability less than 80% were removed. The expanded ion table was summarized by aggregating entries of individual phosphorylation sites, and the best peptide probability and site localization probability for each site were extracted. Total spectral counts were calculated as the sum of all PSMs (peptide spectrum matches) identifying specific phosphorylation site, excluding LC-MS runs with PRM measurements.

### MS data analysis of PRM measurements for phosphorylation site quantification

LC-MS runs with PRM measurements were analyzed in Skyline (22.2.0.351) ^70^. A list containing the 33 peptides targeted by PRM and the raw LC-MS files were imported into Skyline. All peptides and their transitions were validated manually based on retention time, relative ion intensities, and mass accuracy. Extracted ion chromatograms (XICs) were generated for the product ions of all selected peptides, and peak areas were exported from Skyline. The intensity of each peptide was calculated by summing the XICs of the three most intense, interference-free transitions. Subsequently, peptide intensities across different samples were normalized using four reference peptides. First, the sum of the reference peptide intensities was calculated for each sample. Next, these summed intensities were divided by the average across all samples to derive normalization factors. Finally, peptide intensities of each sample were divided by the respective normalization factor. To ensure reliable data, peptides with incomplete quantification or exhibiting an average coefficient of variation exceeding 2 between the two replicates were excluded from further analysis. The intensities of modified peptides covering the same phosphorylated protein site were summed to create site-level intensities. The intensities of the corresponding unmodified peptide counterparts of each site were also summarized to create site level counter intensities. Estimated site occupancy was calculated as “Site intensity” / (“Site intensity” + “Counter intensity”) * 100. For plotting, site-level intensities were log2 transformed and normalized to the average intensity of the respective control samples. The mass spectrometry proteomics data have been deposited to the ProteomeXchange Consortium (http://proteomecentral.proteomexchange.org) via the PRIDE partner repository ^71^ with the dataset identifier PXD045761).

### Protein Expression and Purification

Full-length IGF2BP1 constructs were cloned into a pGEX-6P-2 vector containing an N-terminal GST-tag and a 3C cleavage site. These proteins were expressed in Rosetta (DE3) cells grown to an OD of ∼0.7 and induced with 400 µM IPTG. Cells were grown over night at 20°C, resuspended in GST lysis buffer (25 mM HEPES pH 7.2, 1 M NaCl, 5 % glycerol, 2 mM DTT, 0.5 mM PMSF, 2 mM EDTA), pelleted and frozen in liquid nitrogen.

mCherry-IGF2BP1 constructs were cloned into a pET-47b vector with an N-terminal mCherry- and a C-terminal Deca-His-tag with a 3C cleavage site and expressed in Rosetta (DE3) cells. Shorter IGF2BP1 proteins (KH1/4, KH1/2, KH3/4) were cloned into a pET-47b vector with an N-terminal Hexa -His-Tag and expressed in BL21 cells. His-tagged proteins were induced with 800 µM IPTG and cells were resuspended in His lysis buffer (25 mM HEPES pH 7.2, 1 M NaCl, 5 % glycerol, 5 mM beta-mercaptoethanol, 0.5 mM PMSF, 20 mM Imidazole).

Linker peptides were cloned into a pET-21 vector with an N-terminal His- and SUMO-tag. These proteins were expressed in BL21 cells by growing them in to an OD of ∼0.7, induction with 1mM IPTG and incubation for 3.5 h at 25 °C. The cells were than resuspended in phosphate lysis buffer (20 mM phosphate buffer pH 7.2, 1 M NaCl, 5 % glycerol, 5 mM beta-mercaptoethanol, 0.5 mM PMSF, 20 mM Imidazole). The obtain ^15^N labelled linker proteins, the cells were grown in M9 minimal medium supplemented with ^15^NH_4_Cl. To get ^13^C-^15^N-labelled proteins, glucose was replaced with ^13^C-glucose.

Proteins were purified by resuspending the pellet in lysis buffer and lysing the cells in a EmulsiFlex C3 homogenizer. After pelleting non-soluble cell debris by centrifugating the lysate at 40000 rcf for 30 min at 4 °C, the supernatant was loaded onto the respective affinity column. Full-length proteins were loaded onto two serially connected GST HiTrap HP 5 mL columns, washed with GST wash buffer (25 mM HEPES pH 7.2, 500 mM NaCl, 5 % glycerol, 2 mM DTT, 0.5 mM PMSF, 2 mM EDTA), and eluted with a gradient of 20 mM reduced glutathione in GST wash buffer. His-tagged or His-SUMO-tagged protein were purified on a HisTrap HP5 mL column by washing them with His wash buffer (25 mM HEPES pH 7.2, 500 M NaCl, 5 % glycerol, 5 mM beta-mercaptoethanol, 0.5 mM PMSF, 20 mM Imidazole) and eluted with a gradient of 1 M Imidazole in wash buffer. Bound nucleotides were removed by loading the proteins onto a Heparin HiTrap HP 5 mL column after diluting the NaCl concentration to 100 mM (with wash buffer without NaCl), washing them with wash buffer (25 mM HEPES pH 7.2, 100 mM NaCl, 5 % glycerol, 2 mM DTT, 0.5 mM PMSF, 2 mM EDTA, which was excluded when purifying His-tagged proteins) and eluting them with a gradient of 1 M NaCl in wash buffer. The GST- and His-buffer was cleaved off by incubating the eluted protein with GST-3C or His-3C overnight at 4 °C. For cleaving off the SUMO-tag, proteins were incubated with SENP. The respective tag and the 3C protease were extracted by running the cleaved protein over a GST HiTrap 5 mL or HisTrap HP 5 mL column by using the same buffers as in the affinity purification. For KH1/2 the His-tag was not removed and thus the overnight incubation with protease and the negative His-affinity purification was omitted. The flow through was pooled and further purified via size exclusion chromatography (SEC). Full-length and KH1/2, KH1/4, KH3/4 (no DTT in the buffer) and all respective constructs were eluted with SEC buffer (25 mM HEPES, 150 mM NaCl, 2 mM DTT). Linker constructs were eluted in phosphate SEC buffer (20 mM phosphate buffer pH 7.2, 150 mM NaCl). Protein concentration was measured by using a Nanodrop (full length proteins, linker) or BCA assay (shorter constructs).

### *in vitro* transcription and RNA purification

The RNAs XBP1 200 nt, EIF2A 200 nt, XBP1 Part A and Part B were produced by adding a T7 promoter sequence (TAATACGACTCACTATAGGG) and using the HiScribe T7 RNA Synthesis Kit (NEB E2040S). Template DNA was digested by adding DNase I and incubation for 15 min at 37 °C. RNA was denatured by boiling it for 5 min in denaturing buffer (10m urea, 1mM EDTA, 0.1% SDS, 0.5 mg/mL xylene cyanol, 0.5 mg/mL bromophenol blue) and run on a 6 % acrylamide, 10M urea TBE (89 mM Tris, 89 mM borate, 2 mM EDTA) denaturing gel in 1x TBE buffer for 2 h at 100 V. The gel was stained with SYBR Gold and the bands containing the RNA with the correct size cut out. The RNA was extracted by crushing the gels with a pestle and shaking in RNase-free H_2_O with 1x SUPERaseIn for 1 h at room temperature. The gel pieces were separated by centrifugation in Spin-X filter tubes for 5 min at 20000 rcf. The RNA was further purified via Phenol-Chloroform extraction by mixing the RNA sample first with a 1:1 volume of phenol, vortexing and centrifugation for 2 min at 20000 rcf then taking off the supernatant and mixing it with a 1:1 volume of chloroform, vortexing and centrifugation for 1 min at 20000 rcf. The supernatant is then mixed with a 10: 1 volume of 3M sodium acetate and 1: 1 volume of ice-cold isopropanol. After precipitating the RNA overnight at -80 °C, the RNA was washed twice with ice-cold 80% ethanol and centrifugation for 30 min at 20000 rcf and 4 °C, dried for 10 min and resuspended in RNase free H_2_O. The RNA concentration was determined by using a Nanodrop.

### EMSA Assays

To investigate protein-RNA interaction via Electrophoretic Mobility Shift Assays, proteins were thawed and centrifuged for 15 min at 20000 rcf and 4°C. RNA was refolded by heating up to 95 °C and cooling down to 25 °C in steps of 2 °C/min. The protein and RNA was mixed in EMSA buffer (25mM HEPES pH7.3, 150mM NaCl, 5% glycerol, 2mM EDTA, 10µg/mL Heparin, 0.5mM TCEP) to the specified concentrations. All samples were incubated on ice for 30 min. A 5% TBE gel with 5% glycerol was prepared by pre-running it for 30 min at 210 V in 0.5x TBE buffer at 4°C. 10 µL per sample and 0.5 µL ladder (RiboRuler low range SM1831) were loaded onto the gel and run for 30 min at 210 V in 0.5x TBE buffer at 4 °C. RNA was stained with SYBR Gold and the gel was imaged with BioRad ChemiDoc or iBright CL1500 and quantified with the corresponding software supplied by the manufacturer. RNA binding was determined by using mean intensities and defining the sample with 0 nM protein as 0 and the gel background as 1 on the y-axis. K_D_’s were calculated in GraphPad Prism by using a dose response curve to fit the data.

### Fluorescence Anisotropy Assays

Proteins were thawed and centrifuged for 15 min at 20000 rcf and 4°C. To be able to measure fluorescence anisotropy the corresponding RNAs were obtained from IDTDNA with a 5’ fluorescein tag. The sample with the highest concentration was prepared in anisotropy buffer (25mM HEPES 7.3; 150mM NaCl, 0.025% Tween 20, 10µg/mL Heparin, 2mM EDTA, 0.5mM TCEP) with 10 nM of fluorescence labelled RNA. A concentration series was created by diluting the sample 1: 1 with 10 nM RNA in anisotropy buffer. Subsequently, the samples were incubated on ice for 30 min. The fluorescence anisotropy was measured in a cuvette using an Edinburgh Instruments FS5 spectrofluorometer at threes wavelengths (515, 520 and 525 nm) with a slit width of 5 nm at 20°C with an excitation wavelength of 485 nm, a slit width 5 nm and a dwell time of 1 s. The G-factor, which was measured once for every experiment and then used to calculate the anisotropy for each sample in the experiment, was determined in the beginning of the experiment by measuring 10 nM RNA in anisotropy buffer. The fluorescence anisotropy value was calculated by the manufacturer’s software. K_D_’s were calculated in GraphPad Prism by using a dose response curve to fit the data.

### RNP Granule formation

To investigate the formation of RNP granules, proteins were thawed and centrifuged for 15 min at 20000 rcf and 4°C. RNA was refolded by heating up to 95 °C and cooling down to 25 °C in steps of 2 °C/min. The proteins were diluted in RNase free buffer (25 mM HEPES pH 7.3, 150 mM NaCl, 0.5 mM TCEP). RNP granule formation was induced by the addition of the RNA with the respective final concentration. Right after RNA addition and mixing 50 µL of the sample were pipetted into the well of a Greiner sensoplate, black, 96-well, glass bottom plate which was coated with 1% Pluronic F-127 by washing the well with 200 µL H_2_O, then incubation with 1% Pluronic F-127 for 2 h at room temperature, four times washing with 200 µL H_2_O, and once washing with 100 µL buffer. The plate was incubated on the microscope for 90 minutes to allow droplet formation while avoiding moving and disturbing the granules. The wells were imaged with a Zeiss Axio Observer Z1 in bright field mode, EC Plan-Neofluar 100x/1.3 Oil M27, Orca Flash 4.0 LT+ Camera, VIS-LED at 50 % intensity and 20 ms exposure time.

For the quantification of the condensates, 5 % of the respective N-terminal mCherry-labelled IGF2BP construct was used in the samples. To avoid non-specific interaction of the mCherry-labeled proteins with the glass bottom, the wells were coated with 2 mg/mL BSA in addition to 1% Pluronic F-127 by washing the well with 200 µL H_2_O, then incubation with 100 µL 1% Pluronic F-127 for 2 h at room temperature, twice washing with 200 µL H_2_O, incubation 100 µL 2mg/mL BSA in PBS, twice washing with PBS and once washing with 100 µL buffer. RNP granule formation was induced by the addition of the RNA with the respective final concentration. Right after RNA addition and mixing 50 µL of the sample were pipetted into the well. Imaging was carried out at the microscope setting describe above with additional acquisition of fluorescence images with a Colibri 7 lamp using the 567 nm LED module and a Zeiss filter set 90 HE, 80 % intensity and 250 ms exposure time. Four adjacent tiles were imaged to increase to field of view area. Two fields of view were imaged and analysed per well and the experiment was performed in triplicates. To analyse the data, the tiles were stitched together using the Zeiss ZEN Blue software and cropped to 3200 x 3200 pixels to have similar sized images. Condensate quantification was performed in ImageJ. Background was subtracted with a rolling ball radius of 200 and a sliding paraboloid, the “Default Dark” threshold was set and the image converted to a mask, watershed was used to separate very close condensates and finally the particles were analysed with a minimum size of 200, outlines and a summary were created. The Welch’s t test was used to determine statistical significance.

### FRAP of RNP granules

To investigate the dynamics of IGF2BP1 in RNP granules, proteins were thawed and centrifuged for 15 min at 20000 rcf and 4°C. RNA was refolded by heating up to 95 °C and cooling down to 25 °C in steps of 2 °C/min. The proteins were diluted in RNase free buffer (25 mM HEPES pH 7.3, 150 mM NaCl, 0.5 mM TCEP). IGF2BP1 full-length wild-type, S181E and Y396E were mixed with 5 % of the respective mCherry-labeled IGF2BP1 construct. A Greiner sensoplate, black, 96-well, glass bottom plate was coated with 1% Pluronic F-127 by washing the well with 200 µL H2O, then incubation with 1% Pluronic F-127 for 2 h at room temperature, twice washing with 200 µL H_2_O, incubation 100 µL 2mg/mL BSA in PBS, twice washing with PBS and once washing with 100 µL buffer. RNP granule formation was induced by the addition of the RNA with the respective final concentration. Right after RNA addition and mixing 50 µL of the sample were pipetted into the well. The plate was incubated at the microscope for 90 minutes to allow droplet formation and avoiding moving and disturbing the granules. FRAP was performed on a Zeiss Axio Observer equipped with a Yokogawa CSU-X1 Nipkow spinning disk unit, EC Plan-Neofluar 100x/1.30 Oil Iris objective, 561 nm DPSS laser, Visitron controller, ET605/70 emission filter and a pco.edge sCMOS camera. Images were acquired with 20% laser power and 100ms exposure time. FRAP was performed with a time interval of 3 s, 3 images were taken before bleaching, then selected regions were bleached with 30% FRAP-Laser power for 1 ms per pixel in 5 cycles and 361 frames were taken in total per experiment.

### Phase Separation Assay

To determine whether Linker 1 or Linker 2 can phase separate on their own, the proteins were thawed and centrifuged for 15 min at 20000 rcf and 4°C. The proteins were diluted in buffer (25 mM HEPES pH 7.2, 150 mM NaCl). A Greiner sensoplate, black, 96-well, glass bottom plate was coated with 1% Pluronic F-127 by washing the well with 200 µL H2O, then incubation with 1% Pluronic F-127 for 2 h at room temperature, four times washing with 200 µL H2O, and once washing with 100 µL buffer. 25µL protein dilution were pipetted into the well. 25 µL of 30 % PEG 8000 in buffer were added and carefully mixed to induce phase separation. The plate was incubated on the microscope for 60 minutes to allow droplet formation. The wells were imaged with a Zeiss Axio Observer Z1 in bright field mode, EC Plan-Neofluar 100x/1.3 Oil M27, Orca Flash 4.0 LT+ Camera, VIS-LED at 50 % intensity and 20 ms exposure time.

### Turbidity Assay

Proteins, RNA and the plate were prepared as described under “RNP granule formation”. After induction of granule formation by adding RNA, 50 µL of the samples were pipetted into the plates, the plate put into the plate reader and the experiment started immediately. We used a BioTek Synergy H1 plate reader to measure turbidity. At first, the samples were shaken for 5 s to ensure homogenous distribution of the granules, then the turbidity was monitored by measuring the absorbance of the sample at 480 nm 46 times with a time interval of 20 s. The data was analysed with GraphPad Prism. The one-phase association equation was used to quantify condensate formation.

### MD simulations

We performed molecular dynamics (MD) simulations of linker 1 wild-type, linker 1 S181E, linker 2 wild-type, and linker 2 Y396E using the Martini 3 force field with rescaled protein-water interactions ^72^. We set the scaling parameter λ = 1.06. All linkers were modelled as polypeptide chains with no secondary structure (coils) using UCSF Chimera ^73^ and then martinised. Each simulation box (30×30×30 nm^3^) was set up with 33 randomly placed copies of the same polypeptide chain, water and 0.15 M NaCl. We performed an energy minimization using the steepest descent algorithm. Then we equilibrated the system by running a 10 ps-long simulation using a 1 fs time step and restraining the position of protein backbone beads by using harmonic potentials with force-constants of 1000 kJ mol^-1^ nm^-2^. Afterwards, we ran another 2.1 ns without restraints in the NVT ensemble and a final equilibration of 21 ns in the NPT ensemble, in both equilibration steps we used a 30 fs time step ns. After equilibration, we ran the production phase using a 20 fs time step. The temperature in the simulation box was controlled by a velocity rescale thermostat ^74^(reference temperature T_ref = 300K, coupling time constant tau_T = 1 ps). The Parrinello-Rahman barostat [Parrinello, 1981] (reference pressure p_ref = 1 bar; coupling time constant τ_p = 24 ps) was used for the last equilibration step and for the production run. The simulations were performed using the Martini 3.0 forcefield ^75^ and the GROMACS 2020.5 software^76^.

The contact maps for cis-interactions were calculated using the Contact Map Explorer Python analysis package [https://contact-map.readthedocs.io/en/latest/index.html] (version 0.7.0). For each simulation, we calculated all the contacts between all atoms of the same polypeptide at each frame of the trajectory (ignoring atoms of 2 neighboring residues in each direction). A contact is defined between two atoms that are within a distance of 0.45 nm. The contributions from all chains at each frame were summed up in a single matrix and normalized by the number of frames and chains. For the final plots, the results were shown in a matrix where a value of contact frequency p corresponds to each pair of residues, where p is the max value of the contact frequencies computed for every atom in the residue pair.

The radius of gyration probability distributions computed from MD simulations were compared to the Analytical Flory Random Coil (AFRC) distribution ^53^. AFRC is an analytical model of unfolded polypeptides that behave as ideal chains, so it is suitable to be used as a reference. We computed the AFRC counterparts of linker 1, linker 2 and their phosphomimetic mutants using the AFRC model available via Google Colab notebook: (https://colab.research.google.com/drive/1WHw8ous7IgcKd2LKYuJLeBTlkdEYoRAk?usp=sharing).

### SAXS experiments

SAXS samples were prepared at concentrations > 10 mg/ml in a 250 µL volume and each experiment performed in duplicate. All samples were measured at beamline BM29 at the ESRF facility in Grenoble. A Superdex 200 Increase 10/300 GL sizes exclusion column was equilibrated with SAXS buffer (25 mM HEPES, 150 mM NaCl, 0.5 mM TCEP). Samples were applied to the column and run with 0.5 ml/min. 1300 Frames with 0.5 frames per second were acquired. We used the ATSAS 3.1.3 data analysis software for data processing. SEC-coupled SAXS data was analysed with CHROMIXS. 25 frames for buffer and sample were selected and averaged. The buffer subtracted data was analysed and plotted in PRIMUSQT.

### SAXS analyses and EOM calculations

The data were processed with the SAXSQuant software (version 3.9), and de-smeared using the programs GNOM ^77^ and GIFT (PCG software). EOM analysis was performed with the ATSAS 2.5 package (EMBL, Hamburg). EOM calculations were carried out using the EOM program ^51^ and using default settings. A random pool of 100,000 independent structures was generated using the primary sequence and the available structure of IGF2BP1 domains. All disordered regions were randomized. Using the built-in genetic algorithm and using the default settings, a subset of a few independent structures were selected that described the experimental SAXS best and used to prepare the figures showing R_G_/D_max_ distributions.

### NMR Experiments

NMR spectra were recorded on a 600 MHz Bruker Avance Neo 600 spectrometer. ^1^H-^15^N Heteronuclear Single Quantum Coherence (HSQC) experiments were performed by thawing the proteins and centrifuging for 15 min at 20000 rcf and 4°C. The proteins were diluted with 20 mM phosphate buffer pH 7.2, 150 mM NaCl, 10 % D_2_O to the desired concentration. For the titration experiments with the folded domains, His-RRM1-2, His-KH1-2 or His-KH3-4 were prepared as above. The sample with the highest protein concentration was prepared first, measured and then diluted 1: 1 with 50 µM ^15^N-Linker 1 or 25 µM ^15^N-Linker 2 for higher dilution samples. All HSQC experiments with Linker 1 were performed at 15 °C. All experiments with 12xUG RNA or XBP1 10 nt RNA were performed in RNase free buffer with 25 mM HEPES pH 7.3, 150 mM NaCl. Experiments with 15N-labeled KH1-4 and L2-KH3-4 were conducted in 25 mM HEPES pH 7.2, 150 mM NaCl, 2mM DTT. For all His-RRM1-2 experiments 2 mM DTT was used.

The spectra were processed and phased using NMRPipe ^78^ and further analysed with CCPNMR Version 3 ^79^. Chemical shift perturbations were calculated with the following formula:

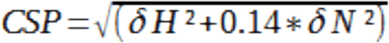

δH and δN are the chemical shift differences compared to the apo protein. Peaks unresolved in the concentrations used for the titration experiments were excluded from the analysis.

Peaks were assigned by obtaining HNCO, HNCACO, HNCACB, HNCOCACB, HNCANNH and HNCOCANNH spectra of 300 µM ^13^C-^15^N-Linker 1 wild-type and 300 µM ^13^C-^15^N-Linker 2 wild-type and the respective HSQC spectra. Peak assignments were performed in CCPNMR. NMR assignments are available in Biological Magnetic Resonance Bank.

### Stress granule reconstitution in cell lysates

Split-GFP tagged IGF2BP1 cells were seeded in a 15cm dish and grown until they reached 90% confluency. Cells were washed with ice cold PBS, and harvested by scraping. Subsequently cells were centrifuged at 3500rpm for 4 minutes. PBS was removed by aspiration, and cell pellet was flash-frozen. In order to lyse the cells, they were thawed and flash-frozen three times. Next, 500ul of lysis buffer, composed of 25mM Tris pH 7, 0.5% NP-40, 1x Protease inhibitor cocktail, 2,5% murine RNAse inhibitor, 100mM NaCl and 2mM DTT, was added. After the addition of the lysis buffer, the cells were further resuspended using a 25G needle 10 times. Next, two centrifugation steps ensued, first at 1500g for 5 minutes, then 16000g for 8 minutes. Supernatants were transferred to a new tube, lysate concentration was measured using a BCA kit (ThermoFisher) and concentration was adjusted to 5mg/ml using the lysate buffer. 20uM of purified G3BP1 were added to induce LLPS followed by a 40 min incubation step. Afterwards, 5uM of mCherry-tagged IGF2BP1 was spiked in, gently mixed and incubated for 20 minutes. The µ-Slide Angiogenesis with ibiTreat 20 ul was used as an imaging vessel. Fluorescence recovery after photobleaching (FRAP) experiments were performed on a Zeiss Axio Observer, using a Plan-Apochromat 63x/1.4 Oil DIC III objective with a Yokogawa CSU-X1 Nipkow spinning disk (50 μm pinholes, spacing 253 μm, 5000 rpm). Imaging was conducted for 3 minutes with 150ms exposure time, with a time interval of 500ms. Three pre-bleach images were taken. 70% of FRAP laser power was used, and a 40% excitation laser.

### Transfection of Packaging Cells

Plasmids used for transfection were purified using an endotoxin-free plasmid kit from Qiagen. Transfections were performed using 1100ng of DNA in total, with 500ng of plasmid of interest, 500 ng pCMVR8.74 (Addgene plasmid # 22036), and 100ng of pCMV-VSV-G (Addgene plasmid # 8454). Supplement free DMEM was used to mix DNA and Polyethylenimine (PEI) in a 1:3 ratio. HEK293T HiEX cell were used as packaging cells. 2*10^5^ cells were seeded in a 6 well plate a day before transfection and grown in a fully supplemented DMEM. The Plasmid mixture containing a transfection reagent was added dropwise onto the cells and they were incubated for 48h.

### Transduction of U2OS (GFP-G3BP1) and HCT116 cells (ΔIGF2BP2, ΔIGF2BP3)

After a 48h incubation, virus was collected from the supernatant with a syringe and sterile filtered. U2OS and HCT116 cells were seeded a day before in fully supplemented medium. The sterile filtered virus was mixed with fully supplemented DMED with 8ug/ml Polybrene at a 1:50 ratio. Cells were grown for 48h up to 15cm dishes. The BD Melody Fluorescence Activated Cell Sorting (FACS) system was used to sort U2OS cells in purity mode, and gated for high and low expressors. The high-expressing population was used for further experiments. HCT116 cells were sorted in a single cell sorting mode, gated for high and low expressors. Three clones were selected for further characterization.

### Western blotting

80% confluent cells were lysed with RIPA buffer (150 mM NaCl, 1% NP-40, 0.5% Sodium deoxycholate, 0.1% SDS, and 25mM TRIS pH 7.4) containing 1x Protease inhibitor (Roche). The protein concentration was determined using a commercially available BCA kit (ThermoFisher). 10 µg of protein containing lysate in sample buffer was denatured at 95°C for 5 minutes. Following denaturing, the samples were loaded onto the 10% sodium dodecyl sulphate (SDS) gel. Proteins were subsequently transferred onto a Nitrocellulose membrane (Amersham) in transfer buffer (25 mM TRIS, 190mM glycine, 20% ethanol) for 110 minutes at 120V. Membranes were stained with Ponceau S and blocked in 5% milk for 1h, or in LI-COR blocking buffer (part number: 927-60001). The primary antibody was diluted in 2.5% milk (Table 1) and incubated overnight at 4°C, or in case of the LI-COR secondary antibody in the blocking buffer. The membrane was washed 5 times with TBST, and the secondary antibody was applied and incubated for 1h, which was diluted in 2.5% milk (Table 1). LI-COR secondary antibody was diluted in the manufacturers antibody diluent (927-65001). After the incubation the membranes were washed 5 times with TBST and an enhanced chemiluminescent (ECL) horse radish peroxidase substrate (ThermoFisher) was added. In case of LI-COR secondary antibodies, after the TBST washes, the membrane was washed three more times with 1x TBS. Membranes were imaged using a BioRad Chemidock, and analyzed using the manufacturers image analysis software (Biorad Image Lab), or using LICOR Odyssey CLx fluorescence imager, and analyzed using Fiji.

| **Antibodies and dilutions used** |  |  |  |
| --- | --- | --- | --- |
| Antibody | Dilution | Manufacturer | Catalog number |
| Anti-IGF2BP1 | 1:2500 | Proteintech | 22803-1-AP |
| Anti-GAPDH | 1:20000 | Proteintech | 10494-1-AP |
| Anti-Rabbit IgG HRP conjugate | 1:20000 | Promega | W401B |
| Anti-Rabbit IgG IRdye 800CW secondary antibody | 1:20000 | LI-COR | 926-32211 |

### Immunofluorescence and Image analyses

20.000 U2OS and HCT116 cells were seeded in an Ibidi µSlide 8 well dish one day before the experiment and grown under the conditions described above. In order to induce the stress granule formation, cells were stressed with 500uM of Sodium arsenite (As) for 30 or 60 minutes. Afterwards cells were washed with PBS and fixed with ice-cold Methanol for 5 minutes. After the fixation the slide was washed 3 more times with PBS and incubated with blocking buffer (5% BSA in TBST) for 1h. In the case of HCT116 cells, the primary ChromoTek RFP-Booster Alexa Fluor 568 (rb2AF568) was incubated overnight. Following the incubation, three washes with PBS ensued and the sample was imaged in PBS using a Zeiss inverse point LSM980 scanning confocal microscope. The objective used was an oil 63x Plan-Apochromat objective with 1.4 NA. The software used for microscope operation is Zeiss ZEN v3.3. The fluorophores were excited with 488nm and 561nm lasers respectively and image analysis was done in Fiji/ImageJ. Cell pose was used in order to get the individual cells in the image, which were saved as an ROI. Cells that touch the image border were excluded, gaussian blur was applied (σ=2), background was subtracted using a rolling ball algorithm with a radius 5, and an auto-threshold “Yen” was applied, after which the particles were analyzed, from sizes 10 pixels to infinity.

### Live cell imaging and FRAP analyses

Live U2OS cells expressing GFP-tagged G3BP1 and mCherry tagged IGF2BP1 were imaged using a Nikon Ti2-E, inverse microscope with a Nikon Perfect Focus System, and a Yokogawa CSU-X1-A1 Nipkow spinning disk (50μm pinholes, spacing 253μm, 5000 rpm). CFI Plan Apo 60x/1.42 Oil, WD 0.15 mm objective was used. Cells were imaged at 37°C and 5% CO_2_. Cells were imaged in either Ibidi µSlide 8 well dish (80826) or Ibidi µdish 35mm (81156). 20000 cells were seeded a day before the experiment in fully supplemented DMEM, and were treated with 500uM Sodium arsenite in Gibco Imaging DMEM (11880-028) for 30 minutes before imaging. Cells were left at 37°C, 5% CO_2_ for 10 minutes to equilibrate before imaging. The fluorescence recovery after photobleaching (FRAP) experiment was conducted with 100% 561 nm and 488 nm laser power for bleaching and with 5ms bleach time per pixel. 5% excitation power was used for the 488 nm laser and 50% for the 561 nm laser during imaging with 150ms of exposure time. The recovery was followed for 8 minutes in a timeseries with the time interval of 3 seconds, with 3 pre-bleach images. Image processing was conducted in Fiji/ImageJ and Excel by selecting the bleached ROI and two ROIs of the similar size with stress granules that were not bleached for bleaching correction. Area without stress granules was also selected in order to conduct background correction. The background is the first to be subtracted, following by the bleach correction, giving us the final values, which are fitted using one phase association in GraphPad PRISM with the formula Y=Y0 + (Plateau-Y0)*(1-exp(-K*x)).

### Immunoprecipitation

HCT116 cells stably expressing mCherry tagged IGF2BP1 were seeded in 15cm dishes and grown until they reached 80% confluency (10^6^ cells). Cells were washed with ice-cold PBS, scraped, centrifuged (3500rpm, 4min) and the pellet flash frozen. On the day of the experiment, the pellets were thawed and 250ul of lysis buffer (consisting of 25mM HEPES pH 7.4, 150mM NaCl, 0.5% NP-40, 0.5mM EDTA, 10% Glycerol, 1x protease inhibitor cocktail (Roche), was added right before usage). Cells were lysed by vortexing for 15 minutes (3 seconds in 2 minute intervals). The lysate was centrifuged at 20.000 rpm for 15 minutes. 50ul of RPF-Trap magnetic agarose (rtma) were used per 15cm dish. Lysates were incubated at 4°C for 2h rotating. The flowthrough was removed by pipetting, while the beads were magnetized using a magnetic rack. The beads were washed five times with wash buffer (25mM HEPES pH 7.4, 150mM NaCl, 0.5mM EDTA, 10% Glycerol). The samples were submitted on-beads for mass-spectrometry analysis at the Max Perutz Labs Mass Spectrometry Facility.

---

**Supp. Figure 1.**
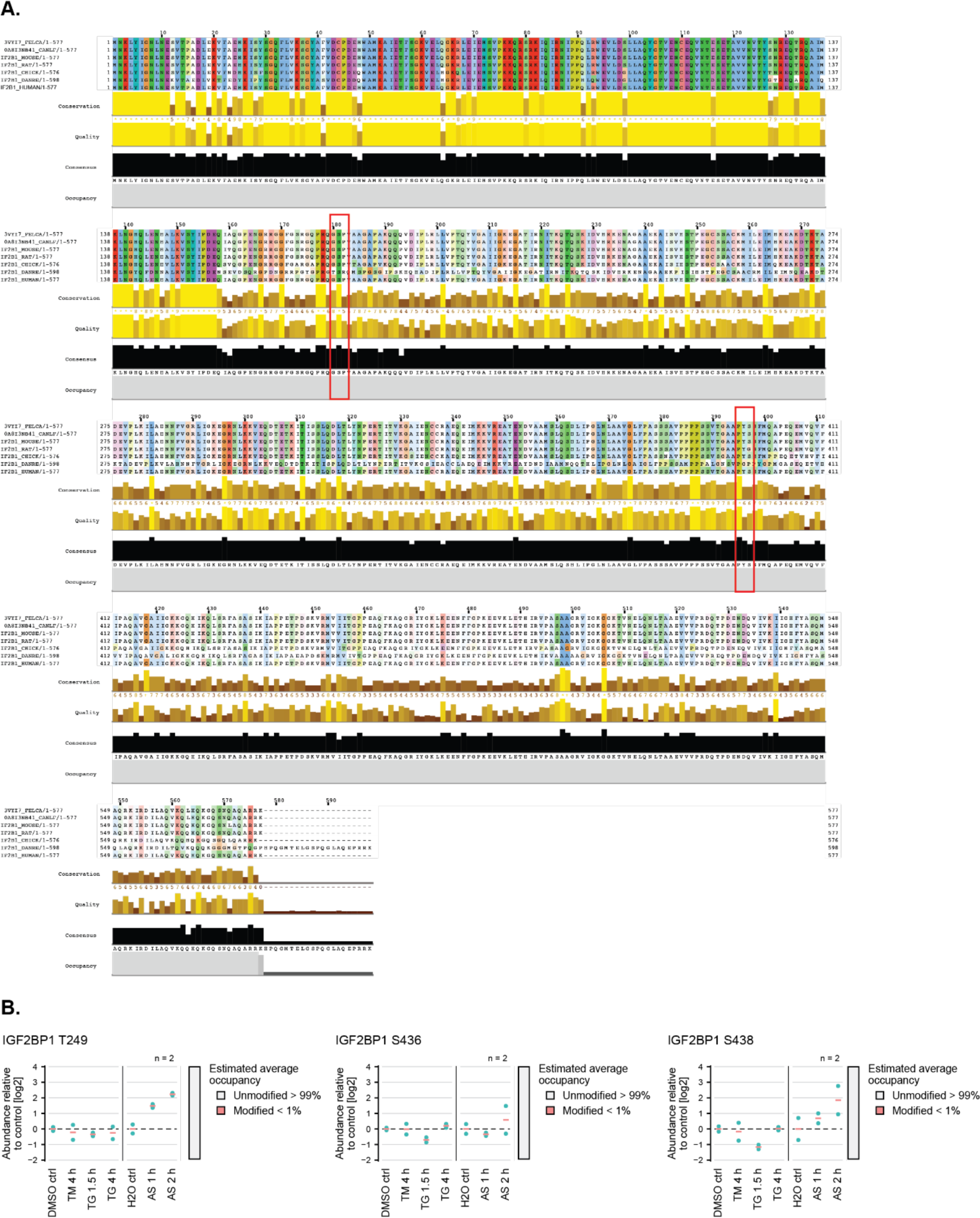
**(A)** Comparison of the IGF2BP1 primary protein sequence from *Felis catus; Canis lupus; Mus musculus; Rattus norvegicus; Gallus gallus; Danio rerio and Homo sapiens.* **(B)** Relative abundance of the indicated IGF2BP1 phosphorylation sites in cells exposed to various forms of proteotoxic stress compared to the control conditions. Tunicamycin (TM) and thapsigargin (TG) induces ER stress, whereas sodium arsenite (AS) leads to oxidative stress. The time-points on the bottom indicate length of exposure to the stress-inducing drug.

**Supp. Figure 2.**
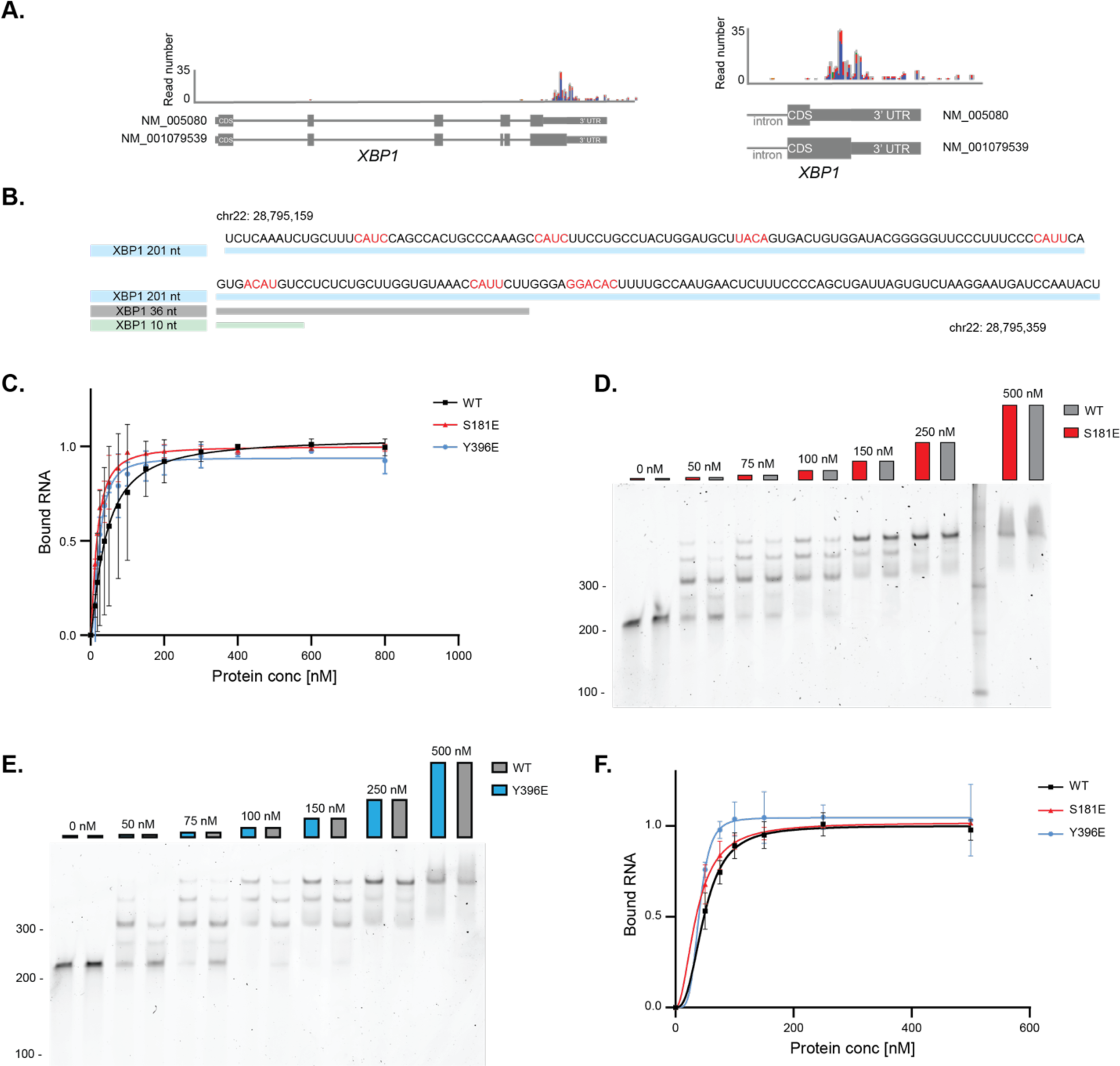
**(A)** IGF2BP1 binding sites in XBP1 identified by analyzing PAR-CLIP data published by Hafner et al. 2010. **(B)** Sequence of the XBP1-derived RNAs: XBP1 201 nt (blue), XBP1 36 nt (grey) and XBP1 10 nt RNA (green). Predicted RNA binding motifs are highlighted in red. **(C)** Quantification of EMSA assays with IGF2BP1 full-length wild-type (black), S181E (red) and Y396E (blue) and XBP1 201 nt RNA from Fig. 2 (A-C) in duplicates. RNA binding is represented by the depletion of free RNA. Mean intensity from 0 nM protein sample was used to define 0 while the mean intensity of gel background represents 1. Dose response equation was used for curve fitting and calculation of K_D_. Error bars represent the standard deviation. **(D) (E)** EMSA assays assessing IGF2BP1 full-length wild-type (black), S181E (red) and Y396E (blue) interaction with EIF2A 200 nt RNA. **(F)** Quantification of EMSA assays of IGF2BP1 full-length wild-type (black), S181E (red) and Y396E (blue) with EIF2A 200 nt RNA from Fig. Supp. 2 (D, E) in duplicates. Dose response equation was used for curve fitting and calculation of K_D_. Error bars represent the standard deviation.

**Supp. Figure 3.**
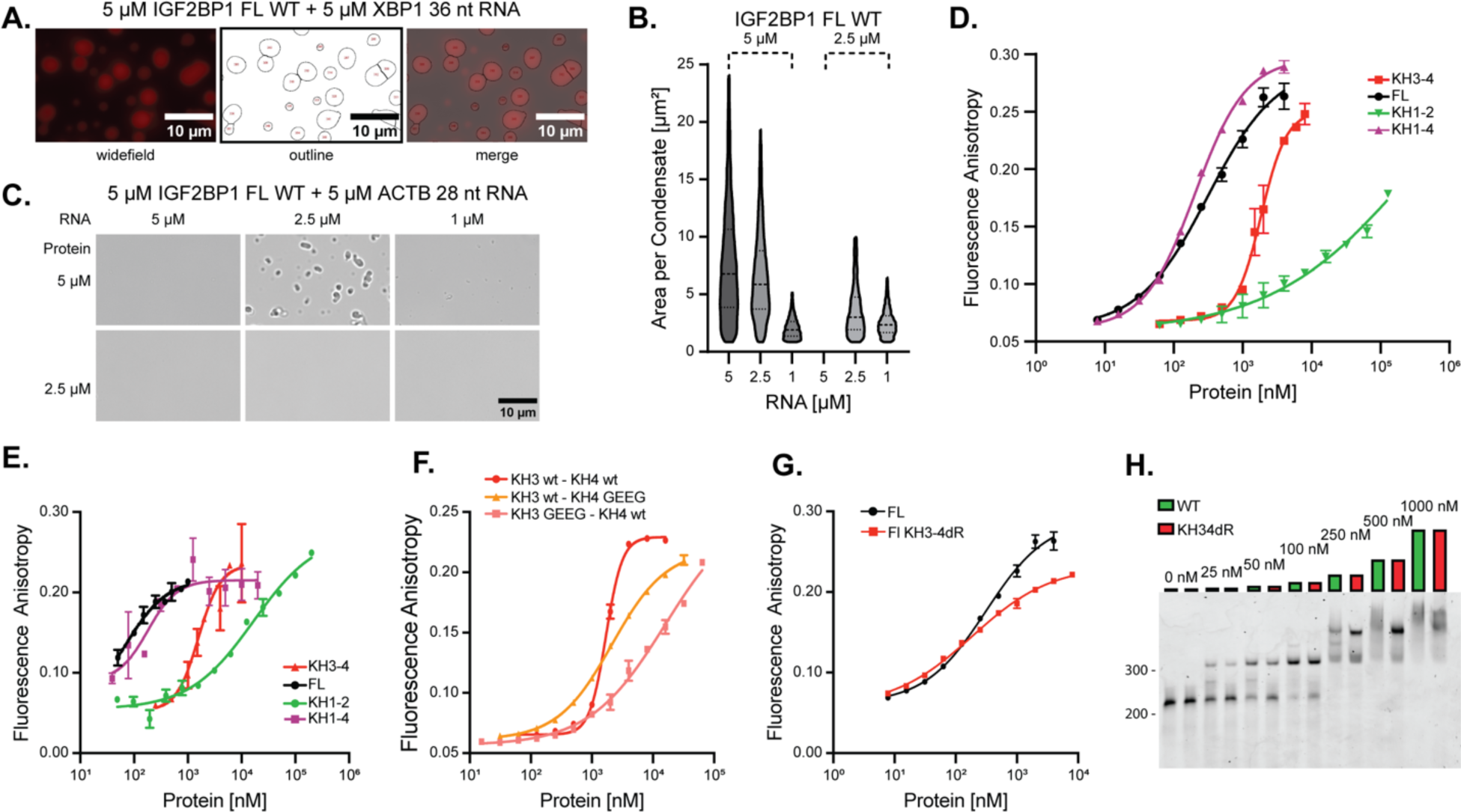
**(A)** Representative images of the quantification of fluorescence microscopy images of condensates formed by 5 µM IGF2BP1 with 5 % mCherry-IGF2BP1 and 5 µM XBP1 36 nt RNA after 90 min of incubation. Outlines created by ImageJ and overlay with the original image. **(B)** Violin plots of the area per condensate after 90 min incubation for IGF2BP1 with XBP1 36 nt RNA at different protein (5 µM and 2.5 µM) and RNA concentrations (x-axis). The dashed line represents the median value, the dotted lines the 25 % and 75 % quartiles. **(C)** RNP granule formation assays of IGF2BP1 with ACTB 28 nt RNA after 90 min of incubation. Scale bar is 10 µm. **(D)** Fluorescence anisotropy assays to assess binding of full-length IGF2BP1 (black), IGF2BP1 KH3-4 (red), IGF2BP1 KH1-2 (green) and IGF2BP1 KH1-4 (purple) binding to 5’-fluorescein labeled XBP1 36 nt RNA. X-axis represented in log-scale. **(E)** Fluorescence anisotropy assays to assess binding of full-length IGF2BP1 (black), IGF2BP1 KH3-4 (red), IGF2BP1 KH1-2 (green) and IGF2BP1 KH1-4 (purple) constructs to 5’-fluorescein labeled ACTB 28 nt RNA. X-axis represented in log-scale. **(F)** Fluorescence anisotropy assays to assess binding of IGF2BP1 KH3-KH4 (red), IGF2BP1 KH3-KH4GEEG mutant (orange) and KH3GEEG-KH4 mutant (pink) with 5’-fluorescein labeled XBP1 36 nt. X-axis represented in log-scale. **(G)** Fluorescence anisotropy assays to assess binding of full-length wild-type IGF2BP1 (black) and full-length IGF2BP1 KH3GEEG-KH4GEEG (KH3-4dR) double mutant (red) with 5’-fluorescein labeled XBP1 36 nt RNA measured by fluorescence anisotropy. X-axis represented in log-scale. **(H)** EMSA assay of full-length wild-type IGF2BP1 (black) and full-length IGF2BP1 KH3GEEG-KH4GEEG (KH3-4dR) mutant (red) with XBP1 201 nt RNA.

**Supp. Figure 4.**
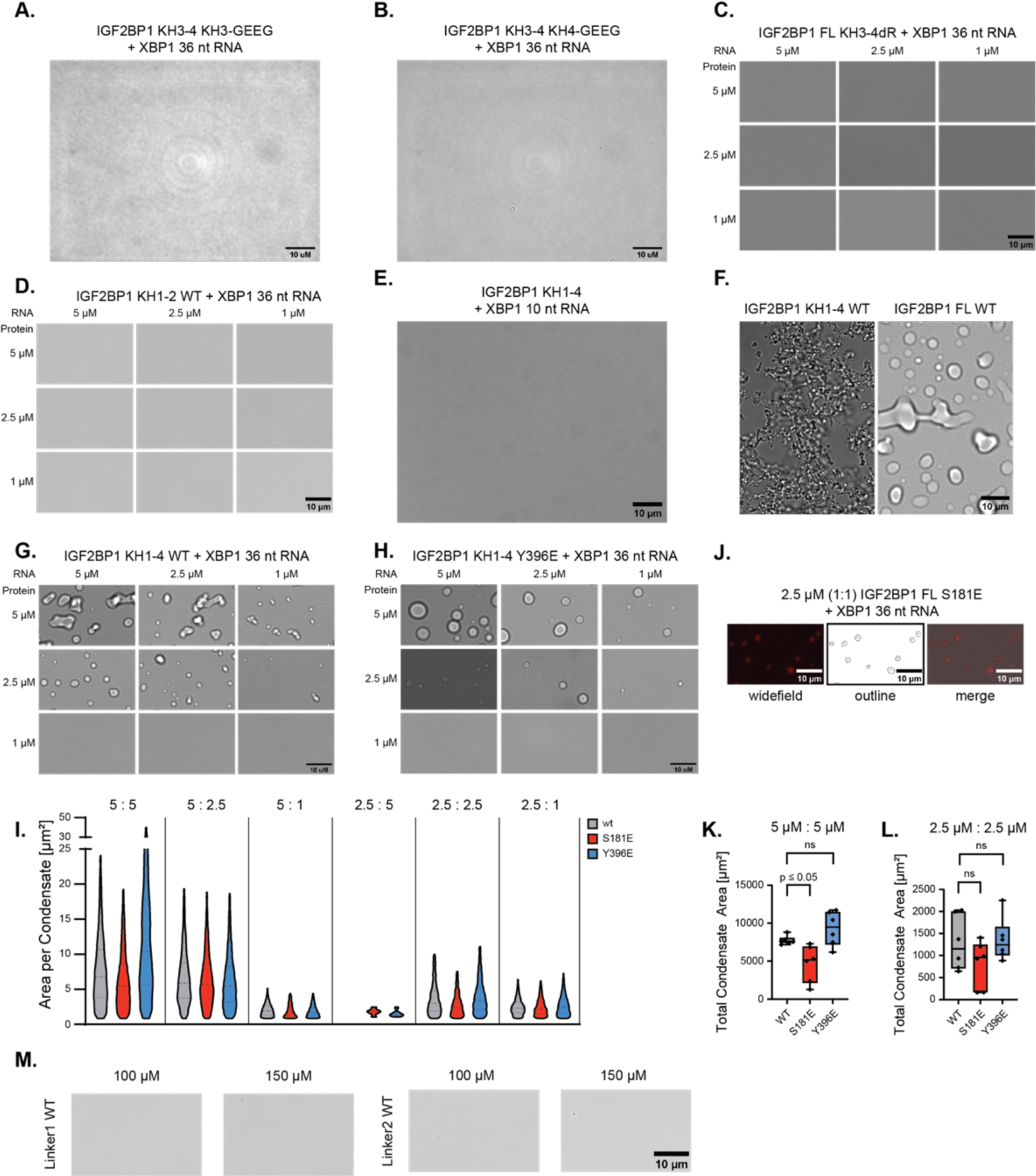
**(A)** RNP granule formation assays of 5 µM IGF2BP1 KH3-4 mutant KH3-GEEG with 5 µM XBP1 36 nt RNA after 90 min of incubation. Scale bar is 10 µm. **(B)** RNP granule formation assays of 5 µM IGF2BP1 KH3-4 mutant KH4-GEEG with 5 µM XBP1 36 nt RNA after 90 min of incubation. Scale bar is 10 µm. **(C)** RNP granule formation assays of IGF2BP1 full-length KH3-4dR (double mutant) with XBP1 36 nt RNA after 90 min of incubation. **(D)** RNP granule formation assays using 6XHis-IGF2BP1 KH1-2 and XBP1 36 nt RNA after 90 min of incubation. Scale bar is 10 µm. **(E)** RNP granule formation assays of 25 µM IGF2BP1 KH1-4 with 25 µM XBP1 10 nt RNA after 30 min of incubation. Scale bar is 10 µm. **(F)** RNP granule formation assay of 5 µM IGF2BP1 KH1-4 in the presence of 5 µM XBP1 36 nt RNA after 90 min incubation in comparison to RNP granules formed by 5 µM full length IGF2BP1 with 5 µM XBP1 36 nt RNA at similar conditions (150 mM NaCl). Scale bar is 10 µm. **(G)** RNP granule formation assays with IGF2BP1 KH1-4 wild-type and XBP1 36 nt RNA at high salt (250 mM NaCl) after 90 min of incubation. Scale bar is 10 µm. **(H)** RNP granule formation assays using IGF2BP1 KH1-4 Y396E and XBP1 36 nt RNA at high salt (250 mM NaCl) after 90 min of incubation. Scale bar is 10 µm. **(I)** Violin plots of the area per condensate after 90 min incubation for full-length IGF2BP1 wild-type, S181E and Y396E with XBP1 36 nt RNA at different protein (5 µM and 2.5 µM) and RNA concentrations (1, 2.5 and 5 µM) and stoichiometries. The first number in the legend shows the protein and the second one is the RNA concentration. The dashed line represents the median value, the dotted lines the upper and lower quartiles. **(J)** Representative images of the quantification of fluorescence microscopy images of condensates formed by 2.5 µM full-length IGF2BP1 S181E in the presence of 5 % mCherry-IGF2BP1 S181E and 2.5 µM XBP1 36 nt RNA after 90 min of incubation. Outlines created by ImageJ and overlaid with the original image. Box plots of total condensate area per field of view in **(K)** 5 µM protein + 5 µM RNA and **(L)** 2.5 µM protein + 2.5 µM RNA. **(M)** Phase separation assay of 100 µM and 150 µM of wild-type linker 1 and wild-type linker 2 in the presence of 15 % PEG 8000 after 60 min incubation. Scale bar is 10 µm.

**Supp. Figure 5.**
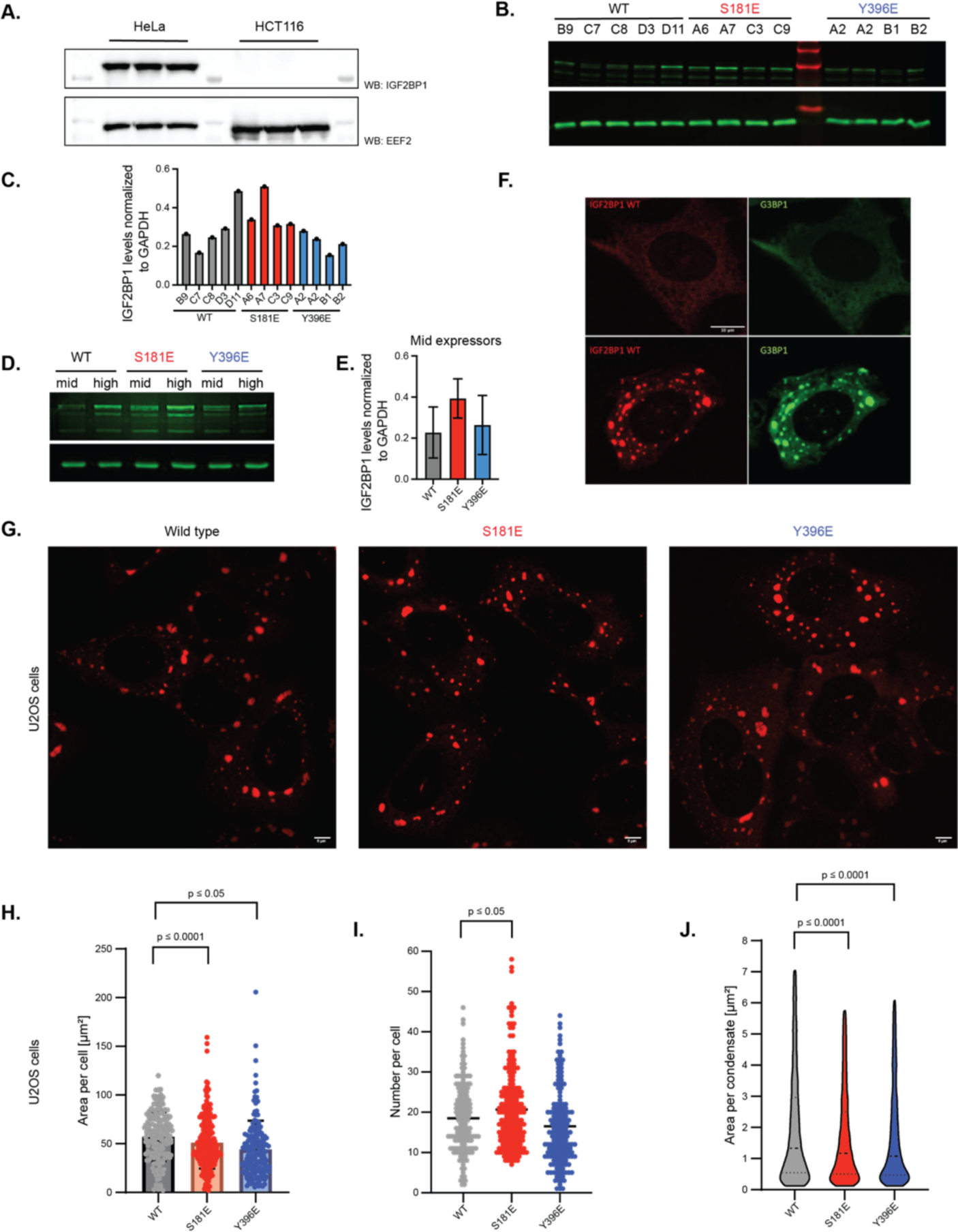
**(A)** Western Blot analyses of IGF2BP1 expression in HeLa and HCT116 cells (Anti-IGF2BP1 antibody, MBL, RN001M; Anti-EEF2 antibody, Proteintech, 20107-1-AP). **(B)** Western Blot analyses of expression level of mCherry-IGF2BP1 wild-type, S181E and Y396E mutants expressed in single clone isolated from HCT116 cells by FACS. The IDs of clones are indicated on the WB. IGF2BP1 is stained by a fluorescently labeled anti-mCherry antibody. **(C)** Quantification of Western Blots from HCT116 cells. The IGF2BP1 band intensities were normalized to bands for the GAPDH control by the LI-COR system for quantification. **(D)** Western Blot analyses of expression level of mCherry-IGF2BP1 wild-type, S181E and Y396E mutants expressed in U2OS cells. Mid and high indicate the expression levels identified by FACS. IGF2BP1 is stained by a fluorescently labeled anti-mCherry antibody. **(E)** Quantification of Western Blots from U2OS cells. The IGF2BP1 bands were normalized to GAPDH control. The LI-COR system was used for quantification. **(F)** Fluorescence microscopy images of fixed U2OS cells expressing mCherry-IGF2BP1 full-length wild-type and GFP-G3BP1 before (top row) and after (bottom row) treatment with 500 µM arsenite for 30 min. **(G)** Representative images of fluorescence microscopy from fixed U2OS cells expressing m-Cherry-IGF2BP1 full-length wild-type, m-Cherry-IGF2BP1 full-length S181E or m-Cherry-IGF2BP1 full-length Y396E stressed with 500 µM arsenite for 60 min. Scale bar is 5 µm. Quantification of condensates in U2OS cells represented as scatter plots: **(H)** total area of condensates per single cell (n = 162 for wild-type, n=230 for S181E, n=162 for Y396E) (bar represents the mean value) **(I)** number of condensates per single cell (bar represents the mean value) and **(J)** area per condensate (the dashed line represents the median value, the dotted lines represent the 25 % and 75 % quartiles).

**Supp. Figure 6.**
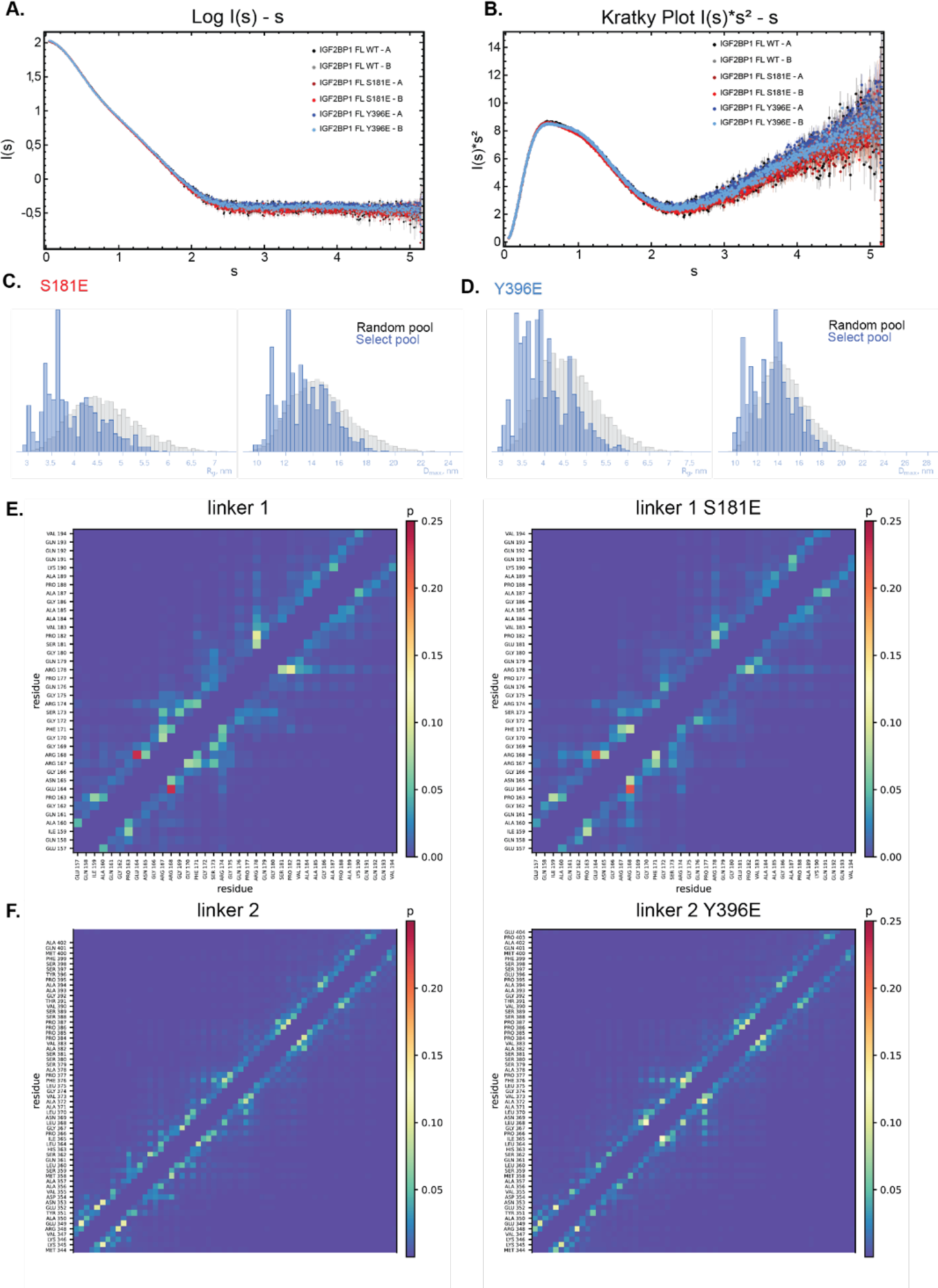
**(A)** Intensity plot I(s) - s of SEC-SAXS data from IGF2BP1 full-length wild-type (black), IGF2BP1 full-length S181E (red), IGF2BP1 full-length Y396E (blue) in duplicates (A, B). Y-axis is represented in log_10_ scale. **(B)** Kratky plot I(s)*s² – s of SEC-SAXS data from IGF2BP1 full-length wild-type (black), IGF2BP1 full-length S181E (red), IGF2BP1 full-length Y396E (blue) in duplicates (A, B). (**C)** Comparison of R_g_ and D_max_ distribution of random conformations of IGF2BP1 S181E and selected pool that best fit the experimental SAXS data based on EOM analyses. **(D)** Comparison of R_g_ and D_max_ distribution of random conformations of IGF2BP1 Y396E and selected pool that best fit to the experimental SAXS data based on EOM analyses. **(E**) Contact map for cis-interactions of linker 1 and its phosphomimetic mutant S181E during MD simulations. **(F)** Contact map for cis-interactions of linker 2 and its phosphomimetic mutant Y396E during MD simulations. Value p measures the frequency of contact formation over the whole MD simulation.

**Supp. Figure 7.**
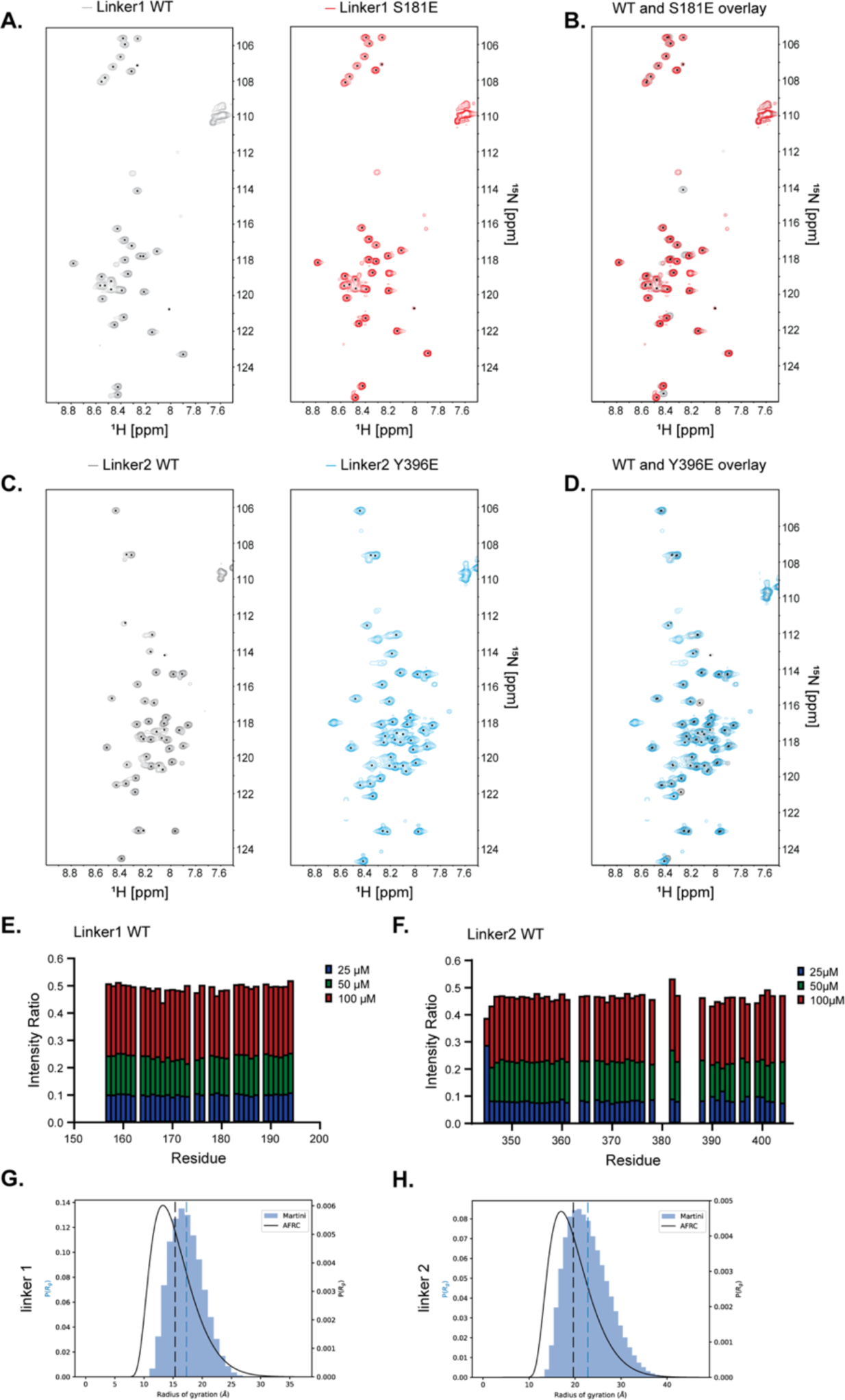
**(A)** HSQC spectra of ^15^N-labeled linker 1 wild-type (grey) and ^15^N-labeled linker 1 S181E (red) at 200 µM. Black dots represent assigned peaks. **(B)** Overlay of the HSQC spectra of linker 1 wild-type and linker 1 S181E (red). **(C)** HSQC spectra of ^15^N-labeled linker 2 wild-type (grey) and ^15^N-labeled Linker 2 Y396E (blue) at 200 µM. **(D)** Overlay of the HSQC spectra of linker 2 wild-type and linker 2 Y396E (blue). Black dots represent assigned peaks. **(E)** Plot of the intensity ratios of 100 µM, 50 µM and 25 µM to 200 µM ^15^N-labeled linker 1 wild-type in HSQC. **(F)**. Plot of the intensity ratios of 100 µM, 50 µM and 25 µM to 200 µM ^15^N-labeled linker 2 wild-type in HSQC. **(G)** Rg probability distribution of linker 1 and **(H)** linker 2 (filled blue steps) overlayed to the probability distribution computed for an ideal polypeptide with the same amino acidic sequence (black line). Dashed lines indicate the mean of the distribution.

**Supp. Figure 8.**
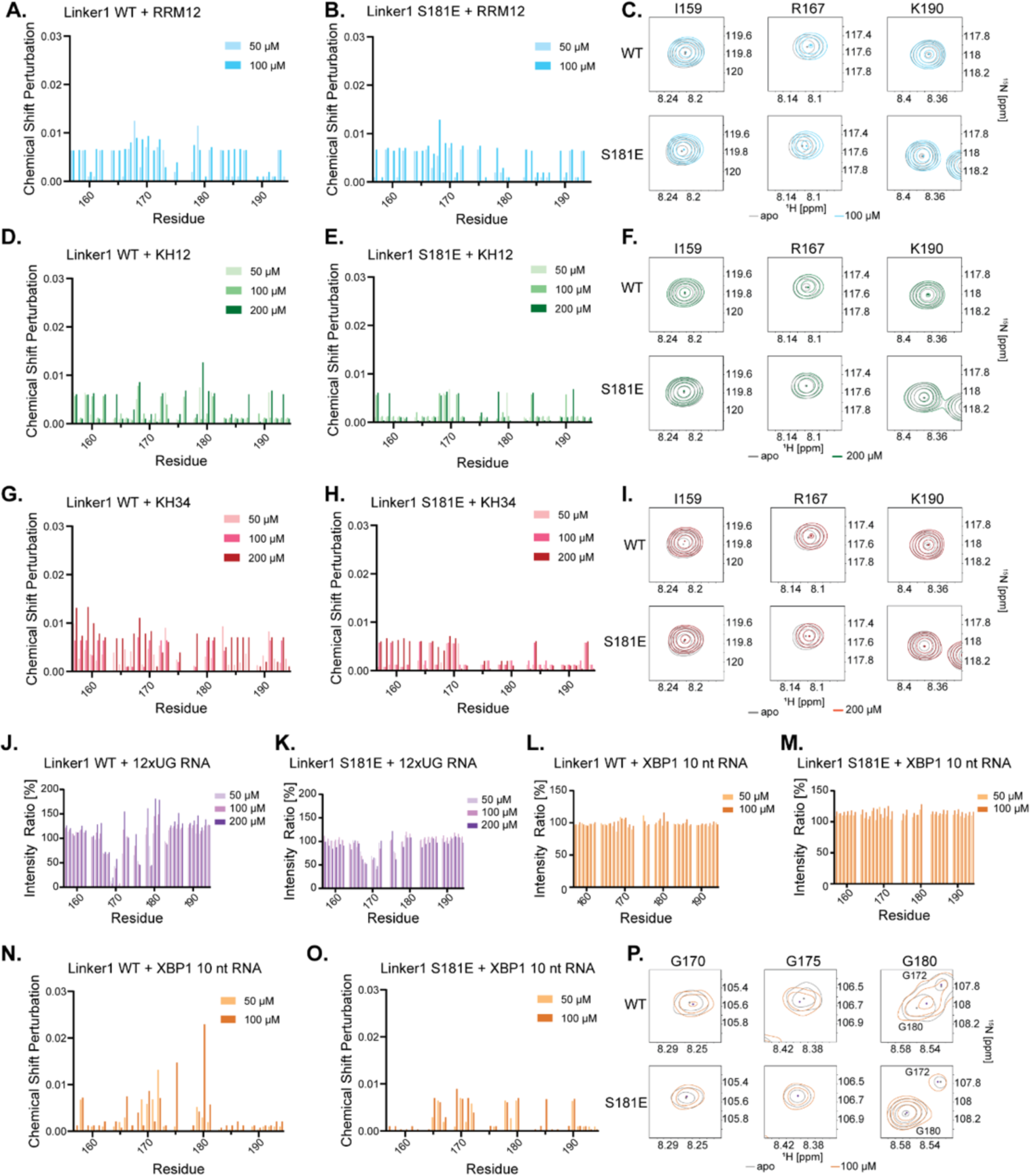
**(A)** CSP analyses of ^15^N-labeled linker 1 wild-type in the absence and presence of different concentrations of RRM1-2 in HSQC experiments. **(B)** CSP analyses of ^15^N-labeled linker 1 S181E with different concentrations of RRM1-2 in HSQC. **(C)** Representative peaks from ^15^N-labeled linker 1 wild-type and S181E alone and with 100 µM RRM1-2 from HSQC spectra. **(D)** CSP analyses of ^15^N-labeled linker 1 wild-type with different concentrations of KH1-2 in HSQC. **(E)** CSP analyses of ^15^N-labeled linker 1 S181E with different concentrations of KH1-2 in HSQC. **(F)** Representative peaks of ^15^N-labeled linker 1 wild-type and S181E alone and with 200 µM KH1-2 from HSQC spectra. **(G)** CSP analyses of ^15^N-labeled linker 1 wild-type with different concentrations of KH3-4 in HSQC. **(H)** CSP analyses of ^15^N-labeled linker 1 S181E with different concentrations of KH3-4 in HSQC. **(I)** Representative peaks of ^15^N-labeled linker 1 wild-type and S181E alone and with 200 µM KH3-4 from HSQC spectra. **(J)** Intensity plots of signals in ^15^N-labeled linker 1 wild-type in the presence of different concentrations of 12xUG RNA in HSQC normalized to signals in the apo spectrum. **(K)** Intensity plots of ^15^N-labeled linker 1 S181E with different concentrations of 12xUG RNA in HSQC normalized to signals in the apo spectrum. **(L)** Intensity plots of ^15^N-labeled linker 1 wild-type with different concentrations of XBP1 10 nt RNA in HSQC normalized to signals in the apo spectrum. **(M)** Intensity plots of ^15^N-labeled Linker 1 S181E with different concentrations of XBP1 10 nt RNA in HSQC normalized to apo spectrum. **(N)** CSP analyses of ^15^N-labeled Linker 1 wild-type with different concentrations of XBP1 10 nt RNA in HSQC. **(O)** CSPs of ^15^N-labeled linker 1 S181E with different concentrations of XBP1 10 nt RNA in HSQC. **(P)** Representative peaks of ^15^N-labeled linker 1 wild-type and S181E alone and with 100 µM of XBP1 10 nt RNA from HSQC spectra.

**Supp. Figure 9.**
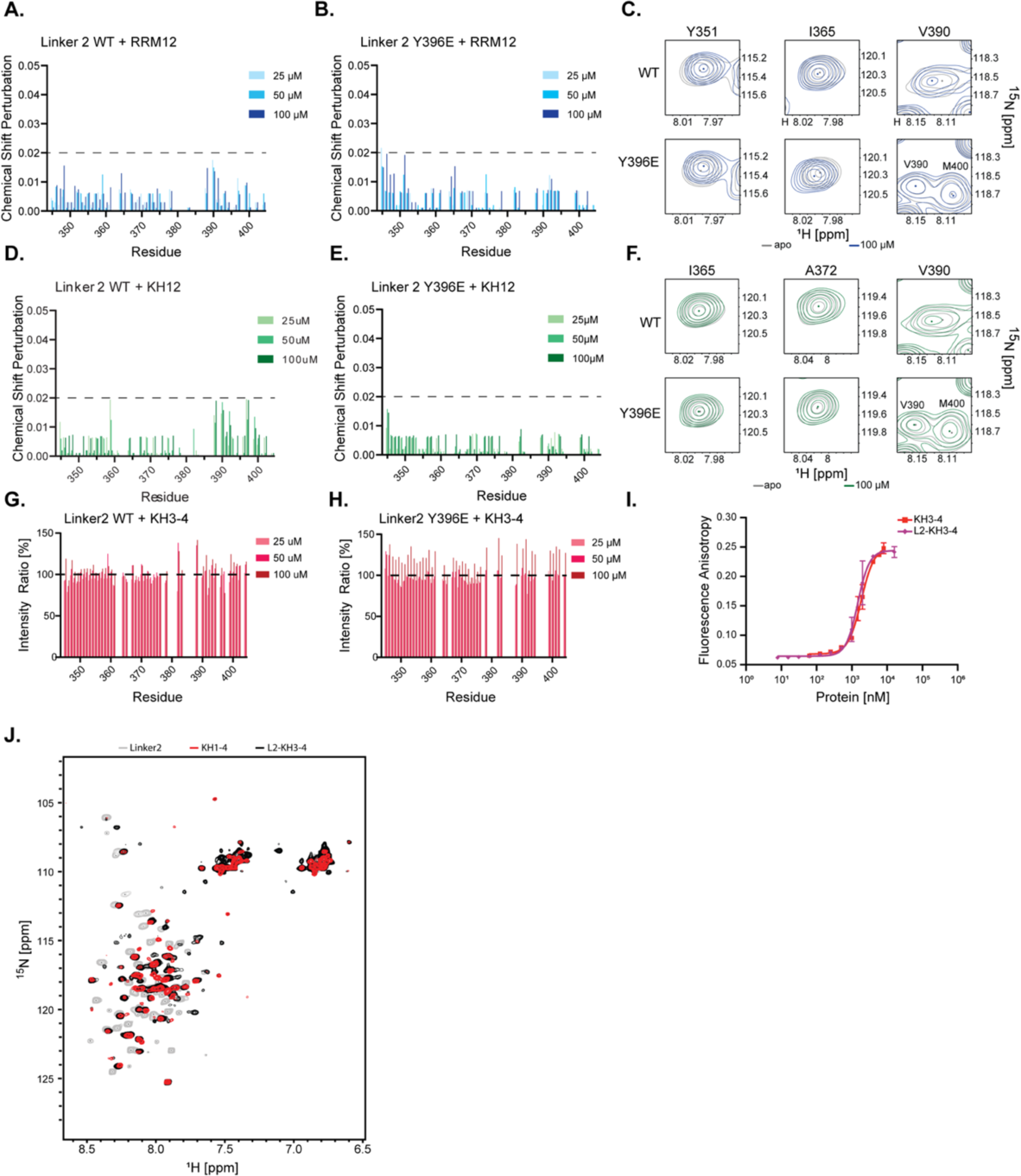
**(A)** CSP analyses of ^15^N-labeled linker 2 wild-type in the absence and presence of various concentrations of RRM1-2 determined by HSQC experiments. **(B)** CSP analyses of ^15^N-labeled linker 2 Y396E with different concentrations of RRM1-2 in HSQC. **(C)** Representative peaks of ^15^N-labeled linker 2 wild-type and Y396E alone and with 100 µM RRM1-2 from HSQC spectra. **(D)** CSPanalyses of ^15^N-labeled linker 2 wild-type with different concentrations of KH1-2 in HSQC. **(E)** CSPanalyses of ^15^N-labeled linker 2 Y396E with different concentrations of KH1-2 in HSQC. **(F)** Representative peaks of ^15^N-labeled linker 2 wild-type and Y396E alone and with 100 µM KH1-2 from HSQC spectra. **(G)** Intensity plots of ^15^N-labeled wild-type linker 2 in the presence of different concentrations of KH3-4 in HSQC normalized to linker 2 signals in the apo spectrum. **(H)** Intensity plots of ^15^N-labeled linker 2 Y396E in the presence of different concentrations of KH3-4 in HSQC normalized to linker 2 Y396E signals in the apo spectrum. **(I)** RNA-protein interactions of IGF2BP1 KH3-4 wild-type (red) and IGF2BP1 Linker 2 KH3-4 wild-type (purple) with 5’-fluorescein labeled XBP1 36 nt RNA measured by fluorescence anisotropy. X-axis represented in log-scale. **(J)** Overlay of HSQC spectra of Linker 2 wild-type (grey), KH1-4 wild-type (red) and Linker 2 KH3-4 wild-type (black).

