## Supplemental Tables for "“IGF2BP1 phosphorylation regulates ribonucleoprotein condensate formation by impairing low-affinity protein and RNA interactions”"

**Table 1:** RNA sequences used in experiments in **Fig.2A-J, 5A-D, Fig. Supp. 2A-F, 3A-H, 4A-L, 8J-P, 9I.** Putative binding sites in **bold** [1-4].

| | **Sequence** |
|---|---|
| ***XBP1* 201 nt** | UCUCAAAUCUGCUUU**CAUC**CAGCCACUGCCCAAAGC**CAUC**UUCCUGCCUACUGGAUGCU**UACA**GUGACUGUGGAUACGGGGGUUCCCUUUCCC**CAUU**CAGUG**ACAU**GUCCUCUCUGCUUGGUGUAAAC**CAUU**CUUGGGA**GGAC**ACUUUUGCCAAUGAACUCUUUCCCCAGCUGAUUAGUGUCUAAGGAAUGAUCCAAUACU |
| ***EIF2A* 200 nt** | AGAAAAUUCAGAAAGA**AACA**GCCCUUCUCCAGGAGCUGGAAGAUUUGGAAUUGGGUAUUUAAAGAUUCACGGAAAGCAAGUUGAUGACCAGAAAUCAGUGCAAA**CACAU**CUUCUGUUAAACC**CAUU**GGUA**UACACA**GAAUAUUCCUGUGCC**CACA**CUUAAUGUCAAUCUAUAAUUUUAAC**CAUU**UAUCCAAGAUUCUACU |
| ***XBP1* 36 nt** | GUG**ACAU**GUCCUCUCUGCUUGGUGUAAAC**CAUU**CUU |
| ***XBP1* 10 nt** | GUG**ACAU**GUC |
| ***ACTB* 28 nt [4]** | ACC**GGAC**UGUUACCAACACC**CACA**CCCC |
| **12xUG** | UGUGUGUGUGUGUGUGUGUGUGUG |


1. Conway, A.E. *et al.* Enhanced CLIP Uncovers IMP Protein-RNA Targets in Human Pluripotent Stem Cells Important for Cell Adhesion and Survival. *Cell Rep* **15**, 666-679 (2016).
2. Schneider, T. *et al.* Combinatorial recognition of clustered RNA elements by the multidomain RNA-binding protein IMP3. *Nat Commun* **10**, 2266 (2019).
3. Hafner, M. *et al.* Transcriptome-wide identification of RNA-binding protein and microRNA target sites by PAR-CLIP. *Cell* **141**, 129-141 (2010).
4. Chao, J.A. *et al.* ZBP1 recognition of beta-actin zipcode induces RNA looping. *Genes Dev* **24**, 148-158 (2010).

**Table2:** $K_D$ values of different full-length IGF2BP1 constructs binding to different RNAs from EMSA and Fluorescence Anisotropy assays in **Fig. 2A-D, Fig. Supp. 2C-F, 3D-H.**

| EMSA | | | |
|---|---|---|---|
| ***XBP1* 201 nt RNA** | $K_D$ [nM] | 95% CI | $R^2$ |
| wild-type | 41.01 | 20.87 to 174.8 | 0.7774 |
| S181E | 17.06 | 14.33 to 20.16 | 0.9727 |
| Y396E | 22.91 | 19.05 to 27.81 | 0.9519 |
| ***EIF2A* 200 nt RNA** | $K_D$ [nM] | 95% CI | $R^2$ |
| wild-type | 48.19 | 42.00 to 53.61 | 0.9686 |
| S181E | 35.42 | 23.25 to 43.23 | 0.9869 |
| Y396E | 40.17 | 14.90 to ??? | 0.9585 |

| Fluorescence Anisotropy | | | | | |
|---|---|---|---|---|---|
| ***XBP1* 36 nt RNA** | $K_D$ [nM] | 95% CI | Hill-Coeff. | 95% CI | $R^2$ |
| wild-type | 311.7 | 258.2 to 390.5 | 0.8640 | 0.7525 to 0.9900 | 0.9932 |
| S181E | 310.1 | 256.3 to 377.5 | 1.049 | 0.8728 to 1.258 | 0.9855 |
| Y396E | 423.3 | 344.8 to 530.7 | 0.9587 | 0.7963 to 1.150 | 0.9862 |
| KH1-4 wild-type | 204.0 | 188.4 to 221.6 | 1.174 | 1.073 to 1.285 | 0.9984 |
| KH1-2 wild-type* | - | - | | | - |
| KH3-4 wild-type | 1850 | 1661 to 2151 | 2.338 | 1.741 to 3.112 | 0.9832 |
| L2-KH3-4 wild-type | 1470 | 1318 to 1640 | 2.620 | 2.042 to 3.392 | 0.9809 |
| Full-length KH3 GEEG KH4 GEEG | 185.8 | 157.4 to 223.2 | 0.6683 | 0.6089 to 0.7328 | 0.9957 |
| KH3 wild-type KH4 GEEG | 2.122 | 1.980 to 2.286 | 0.9702 | 0.9142 to 1.029 | 0.9988 |
| KH3 GEEG KH4 wild-type | 15.69 | 9.738 to 33.85 | 0.6979 | 0.5827 to 0.8249 | 0.9906 |
| ***ACTB* 28 nt RNA** | $K_D$ [nM] | 95% CI | Hill-Coeff. | 95% CI | $R^2$ |
| Full-length wild-type | 68.55 | 56.88 to 87.17 | 1.116 | 0.7966 to 1.471 | 0.9818 |
| KH1-4 wild-type | 245.4 | 24.77 to 374.4 | 3.599 | 0.7449 to ??? | 0.8236 |
| KH1-2 wild-type | 16670 | 10009 to 44411 | 0.7285 | 0.5232 to 0.9849 | 0.9793 |
| KH3-4 wild-type | 1574 | 1175 to 2575 | 2.109 | 1.024 to 4.634 | 0.9285 |

*not enough data points for a reasonable fit

**Table 3:** Area per condensate of IGF2BP1 full-length constructs with XBP1 36 nt RNA (5 µM protein plus 5 µM RNA) in **Fig. 2I**

| | **Median area per condensate [µm²]** | **25% Percentile** | **75% Percentile** | **Total number of condensates** |
|---|---|---|---|---|
| **IGF2BP1 wild-type** | 7.022 | 3.955 | 11.28 | 5377 |
| **IGF2BP1 S181E** | 5.716 | 3.306 | 9.342 | 3897 |
| **IGF2BP1 Y396E** | 10.790 | 5.758 | 18.52 | 4036 |

**Table 4:** Turbidity curve over 15 min of different IGF2BP1 constructs with XBP1 36 nt RNA based on **Fig. 2J.**

| | $t_{1/2}$ [s] | 95% CI | $OD_{480}$ | 95% CI | $R^2$ |
|---|---|---|---|---|---|
| **IGF2BP1 wild-type** | 167 | 138,4 to 205,7 | 0.06775 | 0,06531 to 0,07082 | 0.6448 |
| **IGF2BP1 S181E** | 251.6 | 171,8 to 431,4 | 0.03505 | 0,03130 to 0,04282 | 0.3264 |
| **IGF2BP1 Y396E** | 115.3 | 97,98 to 136,8 | 0.06358 | 0,06230 to 0,06501 | 0.6495 |

**Table 5:** The analyses of the FRAP curves of different IGF2BP1 constructs in G3BP1 induced SGs in **Fig. 3C,D.**

| | **Dynamic pop. [%]** | **95% CI** | **$t_{1/2}$ [s]** | **95% CI** | **$R^2$** |
|---|---|---|---|---|---|
| **IGF2BP1 wild-type** | 68.81 | 67.35 to 70.46 | 21.58 | 19.44 to 24.09 | 0.5531 |
| **IGF2BP1 S181E** | 62.15 | 60.04 to 64.70 | 37.46 | 33.58 to 42.22 | 0.6744 |
| **IGF2BP1 Y396E** | 74.07 | 73.75 to 74.40 | 15.50 | 15.04 to 15.98 | 0.9079 |

**Table 6:** Area per condensate of IGF2BP1 full-length constructs expressed in HCT116 and U2OS cells based on experiments in **Fig.3 E-H, Fig. Supp. 5G-J.**

| **HCT116 cells** | **Median area per condensate [µm²]** | **25% Percentile** | **75% Percentile** | **Total number of condensates** |
|---|---|---|---|---|
| wild-type | 1.142 | 0.4985 | 2.286 | 5272 |
| S181E | 1.026 | 0.4404 | 2.148 | 6719 |
| Y396E | 1.093 | 0.4336 | 2.248 | 3927 |
| **HCT116 cells** | **Median total area per cell [µm²]** | **25% Percentile** | **75% Percentile** | **Mean number of condensates per cell** |
| wild-type | 10.26 | 6.907 | 13.55 | 5.71 |
| S181E | 10.26 | 6.496 | 14.82 | 7.205 |
| Y396E | 10.15 | 6.811 | 15.22 | 6.291 |
| **U2OS cells** | **Median area per condensate [µm²]** | **25% Percentile** | **75% Percentile** | **Total number of condensates** |
| wild-type | 1.666 | 0.6299 | 4.111 | 2980 |
| S181E | 1.407 | 0.5685 | 3.382 | 4463 |
| Y396E | 1.371 | 0.5190 | 3.580 | 2685 |
| **U2OS cells** | **Median total area per cell [µm²]** | **25% Percentile** | **75% Percentile** | **Mean number of condensates per cell** |
| wild-type | 58.36 | 39.04 | 75.63 | 18.54 |
| S181E | 46.3 | 33.6 | 67.83 | 20.66 |
| Y396E | 38.9 | 23.1 | 58 | 16.52 |

**Table 7:** The radius of gyration values of the linkers and their mutants based on the MD simulations in **Fig. 4C-F, Fig. Supp. 7G, H**.

| | **Method** | **<Rg> (Å)** | **<Ree> (Å)** |
|---|---|---|---|
| **Linker 1 WT** | CALVADOS | 16.3 ± 0.1 | 39.1 ± 0.4 |
| | Martini | 17.3 | - |
| | AFRC | 15.31 | 35.81 |
| **Linker 1 S181E** | CALVADOS | 16.25± 0.09 | 39.0 ± 0.4 |
| | Martini | 17.4 | - |
| | AFRC | 15.32 | 35.8 |
| **Linker 2 WT** | CALVADOS | 22.3 ± 0.1 | 52.7 ± 0.7 |
| | Martini | 22.8 | - |
| | AFRC | 19.63 | 45.76 |
| **Linker 2 Y396E** | CALVADOS | 22.2 ± 0.1 | 52.7 ± 0.6 |
| | Martini | 22.5 | - |
| | AFRC | 19.62 | 45.75 |
